# A change in *cis*-regulatory logic underlying obligate versus facultative muscle multinucleation in chordates

**DOI:** 10.1101/2024.03.06.583753

**Authors:** Christopher J. Johnson, Zheng Zhang, Haifeng Zhang, Renjie Shang, Katarzyna M. Piekarz, Pengpeng Bi, Alberto Stolfi

## Abstract

Vertebrates and tunicates are sister groups that share a common fusogenic factor, Myomaker (Mymk), that drives myoblast fusion and muscle multinucleation. Yet they are divergent in when and where they express Mymk. In vertebrates, all developing skeletal muscles express Mymk and are obligately multinucleated. In tunicates, Mymk is only expressed in post-metamorphic multinucleated muscles, but is absent from mononucleated larval muscles. In this study, we demonstrate that *cis-*regulatory sequence differences in the promoter region of *Mymk* underlie the different spatiotemporal patterns of its transcriptional activation in tunicates and vertebrates. While in vertebrates Myogenic Regulatory Factors (MRFs) like MyoD1 alone are required and sufficient for *Mymk* transcription in all skeletal muscles, we show that transcription of *Mymk* in post-metamorphic muscles of the tunicate *Ciona* requires the combinatorial activity of MRF/MyoD and Early B-Cell Factor (Ebf). This macroevolutionary difference appears to be encoded in *cis,* likely due to the presence of a putative Ebf binding site adjacent to predicted MRF binding sites in the *Ciona Mymk* promoter. We further discuss how *Mymk* and myoblast fusion might have been regulated in the last common ancestor of tunicates and vertebrates, for which we propose two models.

## Introduction

In vertebrates, multinucleated myofibers are formed through fusion of mononucleated myoblasts. Myomaker (Mymk) is a transmembrane protein required for myoblast fusion and muscle multinucleation (Millay et al., 2013). In tunicates, the sister group to vertebrates (Delsuc et al., 2006; Putnam et al., 2008), Mymk is also required for myoblast fusion and muscle multinucleation (Zhang et al., 2022). Mymk from tunicate species such as *Ciona robusta* can rescue cell fusion in *Mymk* CRISPR knockout myoblasts in diverse vertebrate species, suggesting highly conserved function (Zhang et al., 2022). Phylogenomic analyses indicate that *Mymk* (previously referred to as *Tmem8c*) arose in the last common ancestor of tunicates and vertebrates through duplication of an ancestral *Tmem8* gene, and is not found in other invertebrates including cephalochordates (Zhang et al., 2022).

However, unlike mammalian *Mymk*, which is expressed in all skeletal muscles, *Ciona Mymk is* exclusively expressed in the differentiating precursors of multinucleated, post-metamorphic (i.e. juvenile/adult) muscles and not in those of mononucleated larval tail muscles (Zhang et al., 2022). Transcription of *Mymk* in mammalian skeletal myoblasts is carried out by Myogenic Regulatory Factor (MRF) family members, especially MyoD1, which function as the molecular switch for muscle specification and differentiation (Zhang et al., 2020). Most tunicates, including *Ciona*, have a biphasic life cycle transitioning from a swimming larval phase to a sessile filter-feeding adult phase (Karaiskou et al., 2015). Their larvae have muscles in their tail that are specified by the *Ciona* MRF ortholog (Meedel et al., 2007). However, unlike the post-metamorphic muscles of the adult body wall and siphons, they do not express *Mymk* and do not undergo cell fusion or multinucleation (Zhang et al., 2022). We therefore sought to understand the molecular mechanism underlying this muscle subtype- and life cycle stage-specific activation of *Mymk* and myoblast fusion in tunicates.

Here, we describe the *cis*- and *trans-*regulatory bases of *Mymk* expression specifically in the post-metamorphic muscles of *Ciona*. First, we show that, like in vertebrates, MRF is required for *Mymk* activation in *Ciona*. However, in *Ciona* the transcription factor Early B-Cell Factor (Ebf, also known as *Collier/Olf1/EBF* or *COE*) is required in combination with MRF to activate *Mymk* transcription. Mis-expressing Ebf together with MRF in the larval tail and other tissues is sufficient to activate ectopic *Mymk* transcription. We show that these effects are recapitulated even by using human MYOD1 and EBF3 in *Ciona*, while *Ciona* MRF alone is sufficient to activate human *MYMK* in cultured myoblasts. Finally, we identify the likely binding sites for MRF and Ebf in the *Ciona Mymk* promoter, suggesting that differences in *cis* (promoter sequences), not in *trans* (transcription factor protein-coding sequences), are the primary drivers of evolutionary change between facultative and obligate *Mymk* expression and myoblast fusion in the chordates.

## Results

Transcription of *Mymk* in mammalian skeletal myoblasts is carried out by bHLH transcription factors of the MRF family (Zhang et al., 2020), which comprise the major molecular switch for muscle specification and differentiation (Hernández-Hernández et al., 2017). In *Ciona*, MRF is the sole ortholog of human MRF family members MYOD1, MYF5, MYOG, and MRF4, and is necessary and sufficient for myoblast specification in the early embryo (Hernández-Hernández et al., 2017; Meedel et al., 2007; Meedel et al., 1997; Meedel et al., 2002). In tunicates, *MRF* is expressed in larval tail muscles and in the atrial and oral siphon muscles (ASMs and OSMs, respectively) of the post-metamorphic juvenile/adult (Razy-Krajka et al., 2014). Yet *Mymk* is only expressed in the ASMs/OSMs (Zhang et al., 2022), suggesting the regulation of *Mymk* in tunicates is different than that of vertebrates as MRF alone is not sufficient to activate *Mymk* in larval tail muscle (**Figure 1A,B**).

**Figure 1.**
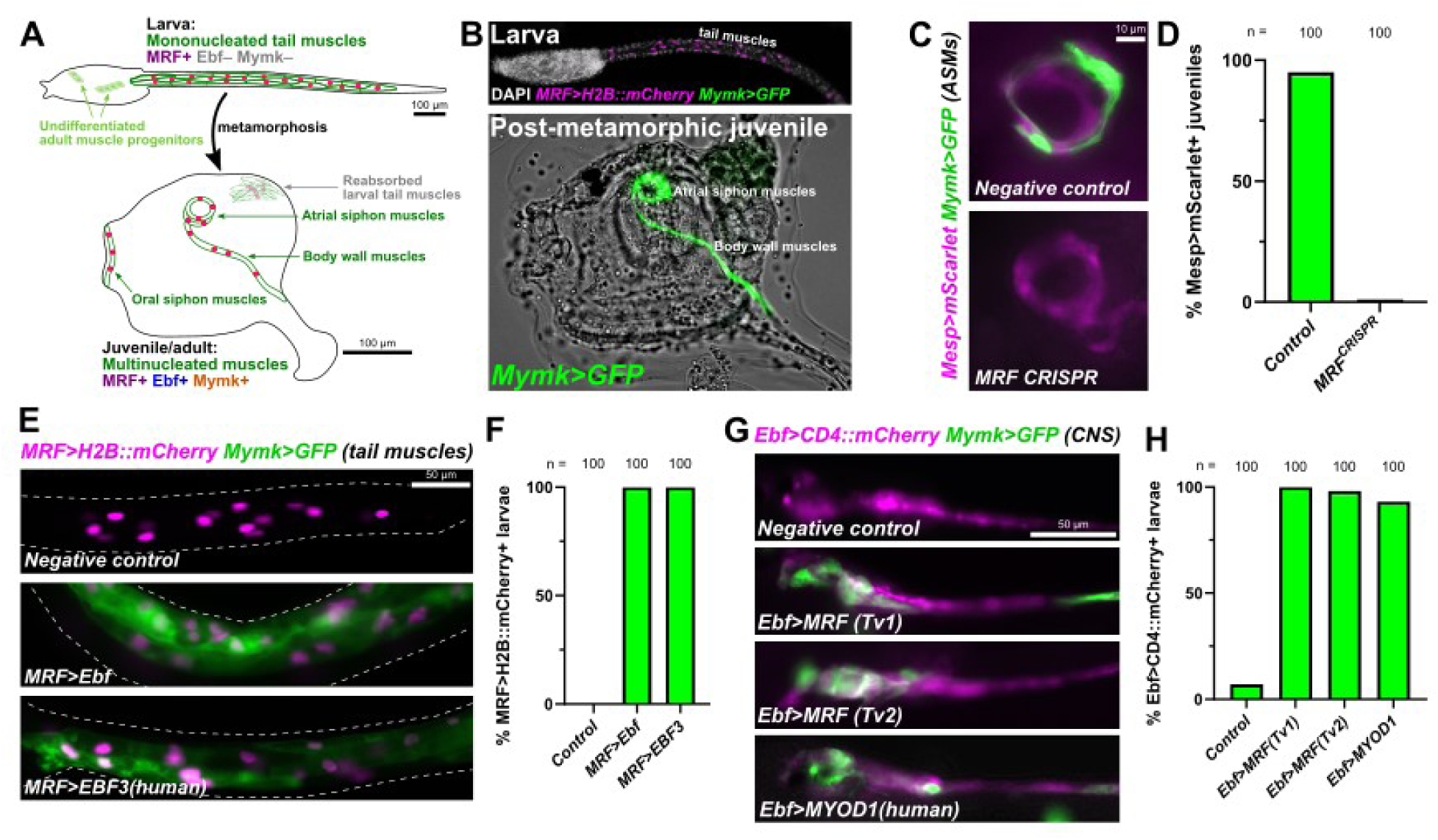
The combination of MRF and Ebf activates *Mymk* expression in *Ciona*. **(A)** Diagram depicting larval or post-metamorphic muscles in the biphasic lifecycle of the tunicate *Ciona robusta.* Based primarily on Razy-Krajka et al. 2014 and Zhang et al. 2022. (**B**) A GFP reporter plasmid containing the entire intergenic region upstream of the *Ciona Mymk* gene (−508/-1 immediately preceding the start codon) is visibly expressed in juvenile/adult muscles at 60 hours post-fertilization (hpf, lower panel) but not in larval tail muscles (21 hpf, upper panel). Juvenile figure re-processed from raw images previously published in Zhang et al. 2022. (**C**) B7.5 lineage-specific CRISPR/Cas9-mediated disruption (using *Mesp>Cas9*) of the *MRF* gene results in loss of *Mymk>GFP* reporter expression (p < 0.0001) in atrial siphon muscles (ASMs). Metamorphosing juveniles fixed and imaged at 46 hpf. Untagged mScarlet reporter used. Negative control juveniles electroporated with *U6>Control* sgRNA vector instead. (**D**) Scoring of data represented in previous panel. (**E**) Ectopically expressing Ebf in MRF+ larval tail muscles (using the *MRF* promoter) results in ectopic activation of *Ciona Mymk* reporter in larvae imaged at 21 hpf. Ectopically expressing human EBF3 in the same cells produces a comparable result. Negative control electroporated with reporter plasmids only. (**F**) Scoring of data represented in previous panel (p < 0.0001 for both Ebf and EBF3). (**G**) Using the *Ebf* promoter to ectopically express either isoform of *Ciona* MRF (Tv1 or Tv2) or human MYOD1 in Ebf+ neural cells results in ectopic activation of *Mymk>GFP* at 16 hpf. Negative control electroporated with *Ebf>lacZ* instead. (**H**) Scoring of data represented in previous panel (p < 0.0001 for all experimental conditions). See text for all experimental details. See **Table S1** for all statistical test details.

Comparing multinucleated post-metamorphic and mononucleated larval muscles, one key molecular difference between the two that we hypothesized might determine the selective regulation of *Mymk* is the expression of Ebf in the former, but not the latter (Stolfi et al., 2010) (**Figure 1A**). Ebf orthologs have been frequently associated with myogenic activity throughout animals. In models such as *Drosophila* and *Xenopus*, Ebf orthologs are upstream of or in parallel to MRF in muscle development (Dubois et al., 2007; Enriquez et al., 2012; Green and Vetter, 2011). In *Ciona*, Ebf specifies post-metamorphic muscle fate (Stolfi et al., 2010; Tolkin and Christiaen, 2016) and activates both *MRF* and ASM-specific gene expression (Razy-Krajka et al., 2014). Therefore, MRF and Ebf were the prime candidates for post-metamorphic muscle-specific activation of *MymK*.

### CRISPR/Cas9-mediated disruption of *MRF* shows it is necessary for *Mymk* expression

To test whether MRF is necessary for *Mymk* expression in post-metamorphic *Ciona* muscles, we targeted the *MRF* locus using tissue-specific CRISPR/Cas9-mediated mutagenesis (Stolfi et al., 2014). We specifically targeted the B7.5 lineage that gives rise to ASMs using the *Mesp* promoter (Davidson et al., 2005) to drive Cas9 expression in this lineage. To target *MRF*, we used a combination of two sgRNAs (*U6>MRF.2* and *U6>MRF.3*) that had been previously designed and validated (Gandhi et al., 2017). We allowed animals to develop into metamorphosing juveniles at 46 hours post-fertilization (hpf) and scored the expression of a previously published *Mymk>GFP* reporter plasmid in *Mesp>mScarlet+* ASMs (**Figure 1C**). In *MRF* CRISPR juveniles, *Mymk>GFP* expression is nearly extinguished, as we observed only 1% of mScarlet+ juveniles showing GFP expression in the *MRF* CRISPR juveniles compared to 95% in the negative control condition (**Figure 1D**). These data strongly suggest that MRF is necessary for *Mymk* transcription in *Ciona* post-metamorphic muscles, just like in vertebrate skeletal muscles.

### Forced co-expression of MRF and Ebf activates ectopic *Mymk* expression

We were not able to utilize the same approach to test the requirement of *Ebf* for *Mymk* expression, as Ebf is required for *MRF* activation in post-metamorphic muscle progenitors (Razy-Krajka et al., 2014), and CRISPR/Cas9-mediated disruption of *Ebf* results in loss of *MRF* expression, converting siphon muscles to heart cell fate (Stolfi et al., 2014). Instead, to test whether the combination of Ebf and MRF is sufficient to activate *Mymk* expression, we forced their combined expression in different larval cells. Outside of being expressed in the progenitors of multinucleated post-metamorphic muscles, *Ebf* is also expressed in the central nervous system of the larva, where it is important for cholinergic gene expression and motor neuron development (Kratsios et al., 2012; Popsuj and Stolfi, 2021). In contrast, *MRF* is expressed in the tail muscles of the larvae, where *Ebf* is not expressed. Therefore we sought to ectopically express Ebf in MRF+ tail muscles, and MRF in Ebf+ neural progenitors.

We first drove ectopic Ebf expression in the larval tail muscles using the extended *MRF* promoter (Zhang et al., 2022). Ectopic expression of EBF in larval tail muscles with *MRF>Ebf* resulted in transcription of the *Mymk* reporter in 100% of successfully transfected larvae compared to 0% in negative control larvae (**Figure 1E,F**). Similarly, we overexpressed MRF in the larval central nervous system using an *Ebf* promoter (Stolfi and Levine, 2011). In this case we tested two different transcript variants (Tv) of MRF, *MRF-Tv1* and *MRF-Tv*2, which differ in the length of the encoded C termini with slightly different functional properties (Izzi et al., 2013). Strong ectopic *Mymk* reporter expression was seen in the larval nervous system with either variant of MRF (MRF-Tv1: 100% *Mymk>GFP*+; MRF-Tv2: 98% *Mymk>GFP*+) (**Figure 1G,H**). In negative control larvae, scattered weak *Mymk>GFP* expression was visible in the nervous system in only 7% of transfected larva. In contrast, overexpression of MRF in the larval tail muscles using an *MRF>MRF-Tv1* construct resulted in *Mymk>GFP* expression in only 8% of larvae, mostly comprising strong expression in certain neurons (presumably Ebf+) that also had leaky *MRF* promoter activity (**Figure S1**) We conclude that increased MRF dose does not adequately replace the function of Ebf. Taken together, our results suggest that MRF and Ebf co-expression is sufficient for *Mymk* activation in *Ciona*.

### Human EBF3 and MYOD1 can replace their *Ciona* homologs

To test whether this MRF-Ebf cooperativity we observe is unique to the *Ciona* proteins, we replaced *Ciona* Ebf and MRF in the above experiments with their human homologs EBF3 and MYOD1. We then assayed whether they can cooperate with endogenous *Ciona* MRF or Ebf and activate the *Ciona Mymk* reporter in the larval tail muscles or central nervous system.

Remarkably, both replacements resulted in strong ectopic *Mymk* reporter expression (**Figure 1E-H**) This implies that the species origin of the proteins themselves does not seem to matter as long as an MRF ortholog and an Ebf ortholog is expressed in the same cell. This suggested that the difference in obligate versus facultative *Mymk* expression is likely due to changes in *cis* (i.e. different binding sites in the *Mymk* promoter) rather than in *trans* (i.e. change to MRF or EBF family proteins) between tunicates and vertebrates.

### *Ciona* MRF alone can activate *MYMK* in human *MYOD1*-CRISPR knockout cells

Using a similar logic from the previous experiment, we introduced *Ciona* MRF (Tv2) and/or Ebf into CRISPR-generated, *MYOD1*-deficient human myoblasts to see whether they (alone or in combination) are sufficient to activate the expression of human *MYMK* (**Figure 2A**).

**Figure 2.**
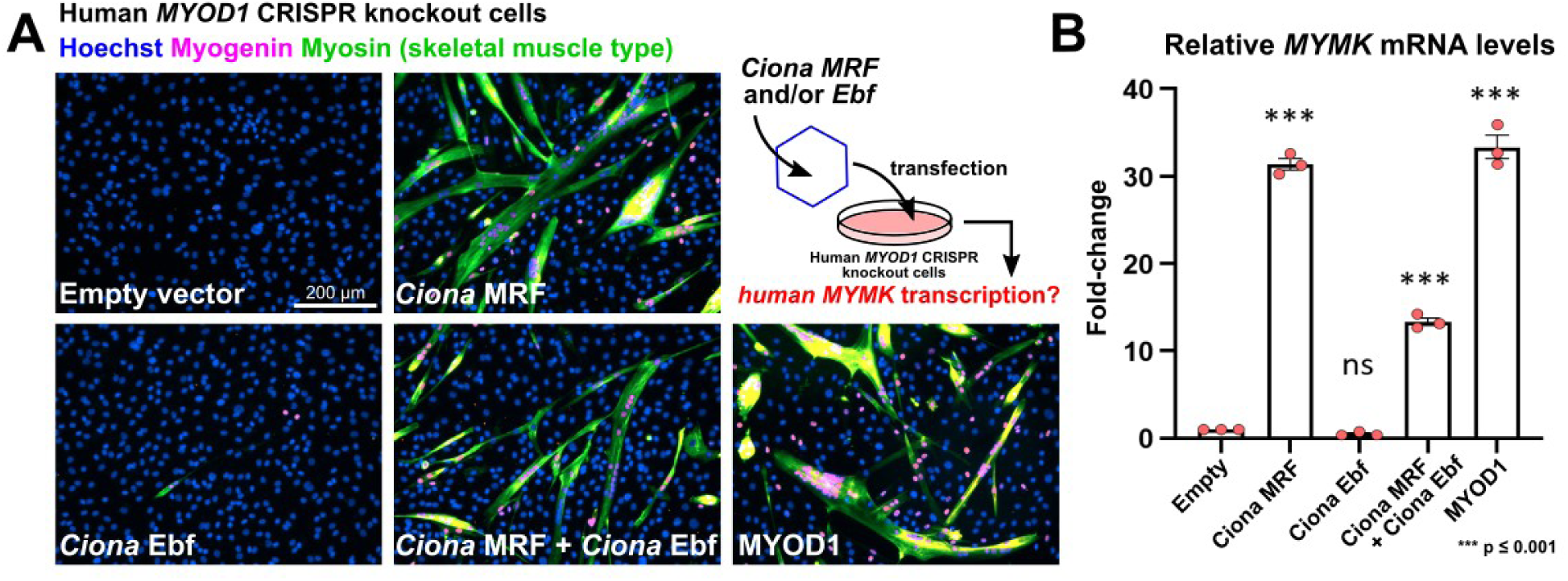
*Ciona* MRF alone can activate transcription of *MYMK* in human cells. **(A)** Representative images of differentiating myoblasts in culture, under different rescue conditions following CRISPR/Cas9-mediated knockout of *MYOD1. Ciona* MRF (Tv2) and Ebf were compared to human MYOD1 for their ability to activate *MYMK,* in combination or solo. Cell nuclei stained with Hoechst (blue), and muscle specification and differentiation visualized with immunostaining for Myogenin (magenta) and skeletal muscle myosin (green). Diagram of rescue experiment in top right panel, with viral vector represented as a hexagon. (**B**) Quantification of *MYMK* mRNA in conditions depicted in previous panel, by qPCR. Experiment performed in triplicate, with statistical significance tested by one-way ANOVA with multiple comparisons to the empty vector condition. See text for experimental details.

Remarkably, transfected *Ciona* MRF alone resulted in nearly identical levels of *MYMK* mRNA expression as human MYOD1 (**Figure 2B**). In contrast, when *Ciona* Ebf was expressed alone, no significant *MYMK* mRNA expression was detected, and Ebf in combination with MRF resulted in a significant reduction of MRF efficacy, appearing to hamper its activation of *MYMK*. These data further support the idea that changes in *cis*, and not in *trans*, underlie the differential requirement of Ebf in activating *Ciona Mymk*.

### RNAseq confirms upregulation of *Mymk* and other post-metamorphic muscle-specific genes by combinatorial activity of MRF and Ebf

Ebf has been established as an important regulator of post-metamorphic muscle fate in *Ciona* (Kaplan et al., 2015; Razy-Krajka et al., 2018; Razy-Krajka et al., 2014; Stolfi et al., 2010; Tolkin and Christiaen, 2016). When we ectopically expressed Ebf in larval tail muscle cells, we observed a striking change in their morphology (**Figure 3A**). Typical larval tail muscles are mononucleated with defined polygonal shapes and cell-cell junctions (Passamaneck et al., 2007), but Ebf expression suppressed these features. Instead, the tail muscle cells became more reminiscent of post-metamorphic siphon and body wall muscles, becoming elongated and myofiber-like, losing their characteristic polygonal shape, and gaining more and smaller-sized nuclei. Although we could not detect any clear instances of tail muscle cells fusing, these observations suggest that the combination of MRF and Ebf might be activating the expression of additional determinants of post-metamorphic muscle-specific morphogenesis.

**Figure 3.**
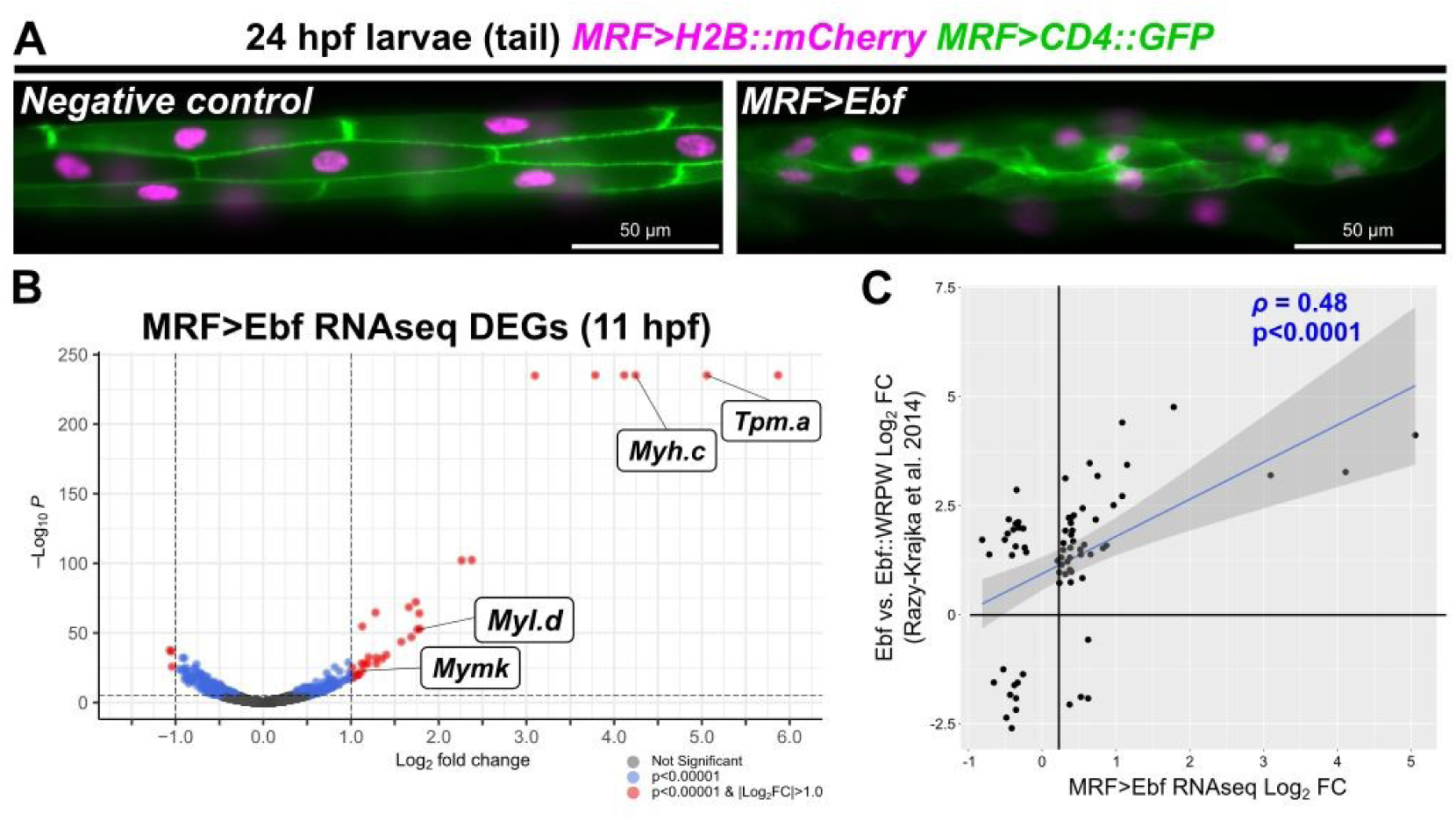
RNAseq analysis of MRF-Ebf targets. **(A)** Morphological phenotype of larval tail muscles altered by overexpression of Ebf (right panel). Note loss of clearly delineated polygonal cell shapes visualized by membrane-bound CD4::GFP (green), compared to the negative control electroporated with the reporter constructs only. (**B**) Volcano plot showing differentially expressed genes (DEGs) detected by bulk RNAseq of whole embryos at 11 hours post-fertilization (hpf), comparing Ebf overexpression (MRF>Ebf) to a negative control condition (MRF>CD4::GFP). *Mymk* and other confirmed post-metamorphic muscle-expressed genes indicated by white boxes: *Tropomyosin.a (Tpm.a), Myosin heavy chain.c (Myh.c,* also known as *MHC3), Myosin light chain.d (Myl.d).* Note the top six genes are flattened at the limit of p-value calculation by the algorithm (see **Table S2** for details and full list of genes). (**C**) Plot comparing our bulk RNAseq data to microarray analysis of DEGs between Foxf>Ebf and Foxf>Ebf::WRPW conditions in FACS-isolated cardiopharyngeal progenitors (CPPs), published in Razy-Krajka et al. 2014 (see reference for original experimental details). Only genes with p<0.05 in both datasets were compared. Rho (ρ) indicates Pearson’s correlation. Dark grey area indicates 95% confidence interval.

To identify other putative targets of MRF-Ebf cooperativity, we performed bulk RNAseq to compare the transcriptomes of wildtype embryos to embryos in which Ebf was overexpressed in the tail muscles. We extracted RNA at 11 hpf from whole embryos that were either transfected with *MRF>Ebf* or, *MRF>CD4::GFP* as a negative control. Using DESeq2 to analyze the resulting RNAseq data, we observed significant upregulation of many genes in the *MRF>Ebf* condition. Of the 16,433 genes detected, 163 were significantly upregulated (p<0.00001, logFC>0), of which 33 showed a logFC greater than 1.0 (**Figure 3B**, **Table S2**). In contrast, 155 genes were significantly downregulated (p<0.00001, logFC<0). Interestingly, high-ranking genes that had both a significant p-value and log fold change ≥ 1 include *Tropomyosin.a (Tpm.a), Col24a-related*, *myosin heavy chain.c (Myh.c),* and *myosin light chain.d (Myl.d)*, which have all been confirmed as upregulated specifically in the ASMs by *in-situ* hybridization (Razy-Krajka et al., 2014). *Mymk* was #25 in this list, confirming that endogenous *Mymk* (and not just the *Mymk>GFP* reporter) is ectopically activated in tail muscles upon Ebf overexpression. This suggests that Ebf is sufficient to partially convert larval tail muscle cells into post-metamorphic, atrial siphon-like muscles.

To further investigate this muscle subtype fate change, we compared our EBF overexpression bulk RNAseq results to a published microarray analysis of Ebf overexpression in the Trunk Ventral Cells (TVCs) that give rise to both heart and ASM progenitors (Razy-Krajka et al., 2014). Indeed, when comparing both data sets, many of the top genes in our list were also significantly upregulated by Ebf overexpression in the TVCs, resulting in a Pearson correlation coefficient (ρ) of 0.48 (**Figure 3C**, **Table S3**). Although there are several genes that show discrepant changes in expression between the two datasets, this may reflect differences in the timing of RNA extraction and territory of Ebf overexpression (tail muscles at 11 hpf vs. TVCs at 21 hpf). Taken together, these data suggest that *Mymk* is just one of several genes that might be preferentially activated in post-metamorphic, multinucleated muscles by a similar MRF-Ebf combinatorial logic in *Ciona*.

### Analyzing candidate MRF and Ebf binding sites in the *Mymk* promoter

Because we suspected vertebrate-tunicate differences in *Mymk* activation to be due primarily to differences in *cis,* we aligned the *Ciona robusta Mymk* promoter to the homologous sequence from the related species *Ciona savignyi* (Satou et al., 2008; Satou et al., 2019; Satou et al., 2022; Vinson et al., 2005) to identify potentially conserved transcription factor binding sites (**Figure 4A**). We also utilized JASPAR (Castro-Mondragon et al., 2022), a predictive binding site search algorithm, to look for putative MRF and Ebf sites (Chaudhary and Skinner, 1999; Treiber et al., 2010). This led us to identifying MRF^-136^ and Ebf^-116^ as conserved, high-scoring candidate binding sites to test (**Figure 4A**, **Table S4**). We made mutations predicted to disrupt MRF or Ebf binding to these putative sites in the *Mymk>GFP* reporter plasmid, and scored the co-expression of these mutant reporters with a wild type (“WT”) *Mymk>mCherry* reporter. When observing juveniles electroporated with the *Mymk>GFP* reporter bearing the MRF^-136^ mutation, it was clear that its activity was significantly reduced, with only 19% of *Mymk(WT)>mCherry+* siphon muscles faintly expressing GFP as well (**Figure 4B**,**C**). As expected, we also observed dramatic reporter expression loss with the Ebf^-116^ mutation (20% GFP expression, although mutating both MRF^-136^ and Ebf^-116^ did not further abolish the residual GFP expression (**Figure 4B**,**C**). In contrast, 100% of juveniles co-expressed wild type GFP and mCherry reporters.

**Figure 4.**
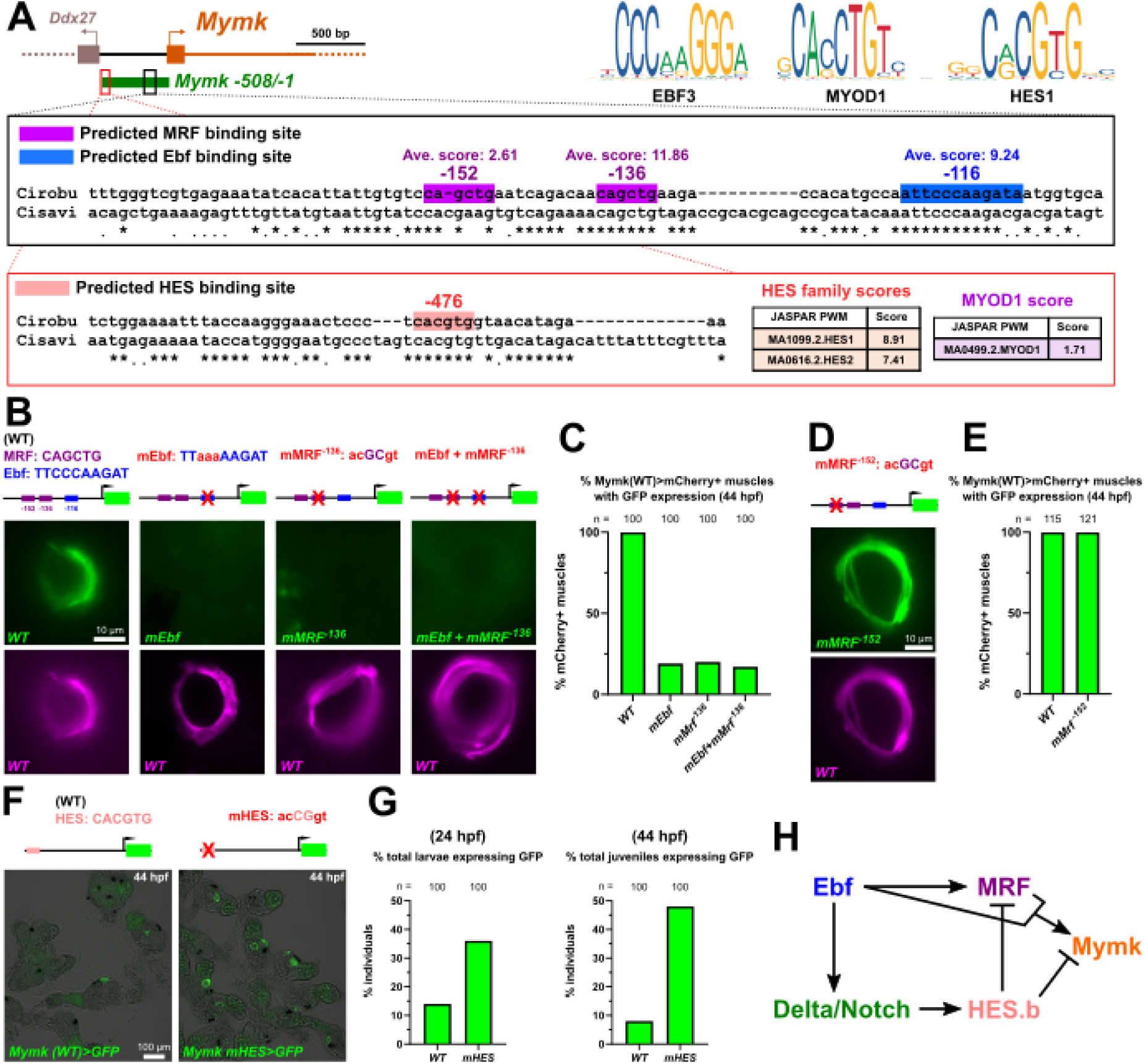
Mutational analysis of predicted binding sites in the *Ciona Mymk* promoter. **(A)** Diagram of *Ciona Mymk* genomic region with predicted binding sites highlighted in insets. Coordinates given as relative to Mymk translational start codon, as transcription start sites are generally unavailable for *Ciona* genes. Conserved basepairs indicated by asterisks under alignment between *C. robusta* and *C. savignyi* orthologous sequences. Top right: position-weight matrices (PWMs) for human orthologs of the major candidate transcription factors analyzed in this study. Bottom inset: Far upstream Ebox (−476) shows greater predicted affinity for HES-family repressors than for MYOD1/MRF activators. Predicted scores obtained from JASPAR. (**B**) Disruptions to predicted binding sites in the *Mymk>GFP* reporter results in significant loss of activity in post-metamorphic siphon muscles, imaged at 44 hours post-fertilization (hpf). All GFP reporters co-electroporated with wild type *Mymk>mCherry* reporter. (**C**) Scoring of data represented in previous panel (p < 0.0001 for all). (**D**) Disrupting the low-scoring, non-conserved MRF^-152^ site does not significantly reduce reporter expression (p > 0.9999), as quantified in (**E**). (**F**) Mutating the predicted HES site at position -476 results in higher frequency of reporter expression. (**G**) Scoring of data represented in previous panel and similarly electroporated larvae at 24 hpf. Total individuals were assayed for GFP reporter expression. Normally, only ∼5-15% of all individuals show *Mymk>GFP* expression, likely due to mosaic uptake/retention of electroporated plasmids. Mutating the HES site boosts this to ∼30-45% (p < 0.0005 at 24 hpf, p < 0.0001 at 44 hpf). (**H**) Gene regulatory network diagram showing proposed regulation of *Ciona Mymk* by Ebf, MRF, and Notch-dependent HES. Regulatory connections between Ebf, MRF, Delta/Notch, and HES.b based on Razy-Krajka et al. 2014. See text for experimental details and **Table S1** for statistical test details.

Similarly, mutation of a nearby poorly conserved, low-scoring predicted MRF site (MRF^-152^) did not significantly reduce reporter activity, suggesting it is not required for activation and likely not a functional MRF binding site (**Figure 4D**,**E**). Taken together, these data suggests that *Ciona Mymk* activation is dependent on closely spaced, conserved MRF and Ebf predicted binding sites in its proximal promoter region.

### Predicted HES binding site represses *Mymk* activation

When examining the *Mymk* promoter for potential transcription factor binding sites, we noticed a conserved Ebox sequence further upstream in the *Mymk* promoter (**Figure 4A**). We initially thought could be an MRF binding site, but JASPAR predictions revealed a much higher score for binding by Hairy Enhancer of Split (HES) transcriptional repressor family members (**Figure 4A**). In Ciona, HES has been shown to mediate Delta/Notch-dependent repression of *MRF* expression and myogenic differentiation in the inner ASM precursor cells, prolonging their undifferentiated, proliferative state (Razy-Krajka et al., 2014). In vertebrates, Delta/Notch signaling also represses MyoD expression and muscle differentiation (Delfini et al., 2000). In chick, HEYL (a HES homolog) binds to the *Mymk* promoter and inhibits its transcription, hinting at a deeply conserved strategy for restricting the onset of *Mymk* expression and fusion in developing myoblasts (Esteves de Lima et al., 2022). When we tested a *Mymk* GFP reporter plasmid carrying a mutation to disrupt this upstream Ebox, we observed a significant increase in frequency of GFP expression compared to the wild type reporter (**Figure 4F**,**G**). Increased GFP expression suggests that this site is most likely bound by a repressor. Our results suggest that the direct repression of *Mymk* transcription by HES repressors (**Figure 4H**) may have been an ancestral trait present in the last common ancestor of tunicates and vertebrates.

### Adding an additional high-quality MRF binding site abolishes the need for MRF-Ebf cooperativity

What might be the exact *cis*-regulatory change that result in the difference observed between tunicate and vertebrate *Mymk* regulation? Unfortunately, reporter constructs made using published human *MYMK* (Zhang et al., 2020) or chicken *Mymk* (Luo et al., 2015) promoters were not expressed at all in *Ciona* tail muscles (**Figure S2**). This was not entirely surprising, given that orthologous promoters are frequently incompatible (i.e. “unintelligible”) even between different tunicate species due to developmental system drift (Lowe and Stolfi, 2018). We therefore focused instead on testing different point mutations in the *Ciona Mymk* promoter that might result in ectopic activation in larval tail muscles.

*Cis*-regulatory logic can be complex with subtle changes in promoter sequences resulting in drastically different activation patterns (Spitz and Furlong, 2012). In tunicates, it has been shown that by making changes to the sequences flanking a given transcription factor binding site one can increase its predicted binding affinity, resulting in higher expression levels or ectopic activation (Farley et al., 2015; Farley et al., 2016; Jindal et al., 2023). To test whether such “optimized” MRF and Ebf binding sites might result in activation of the *Ciona Mymk* reporter by MRF or Ebf alone (without the need for MRF-Ebf cooperation), we manipulated flanking sequences of putative MRF or Ebf binding sites, resulting in higher binding affinity scores predicted by JASPAR (**Figure 5A,C, Figure S3**).

**Figure 5.**
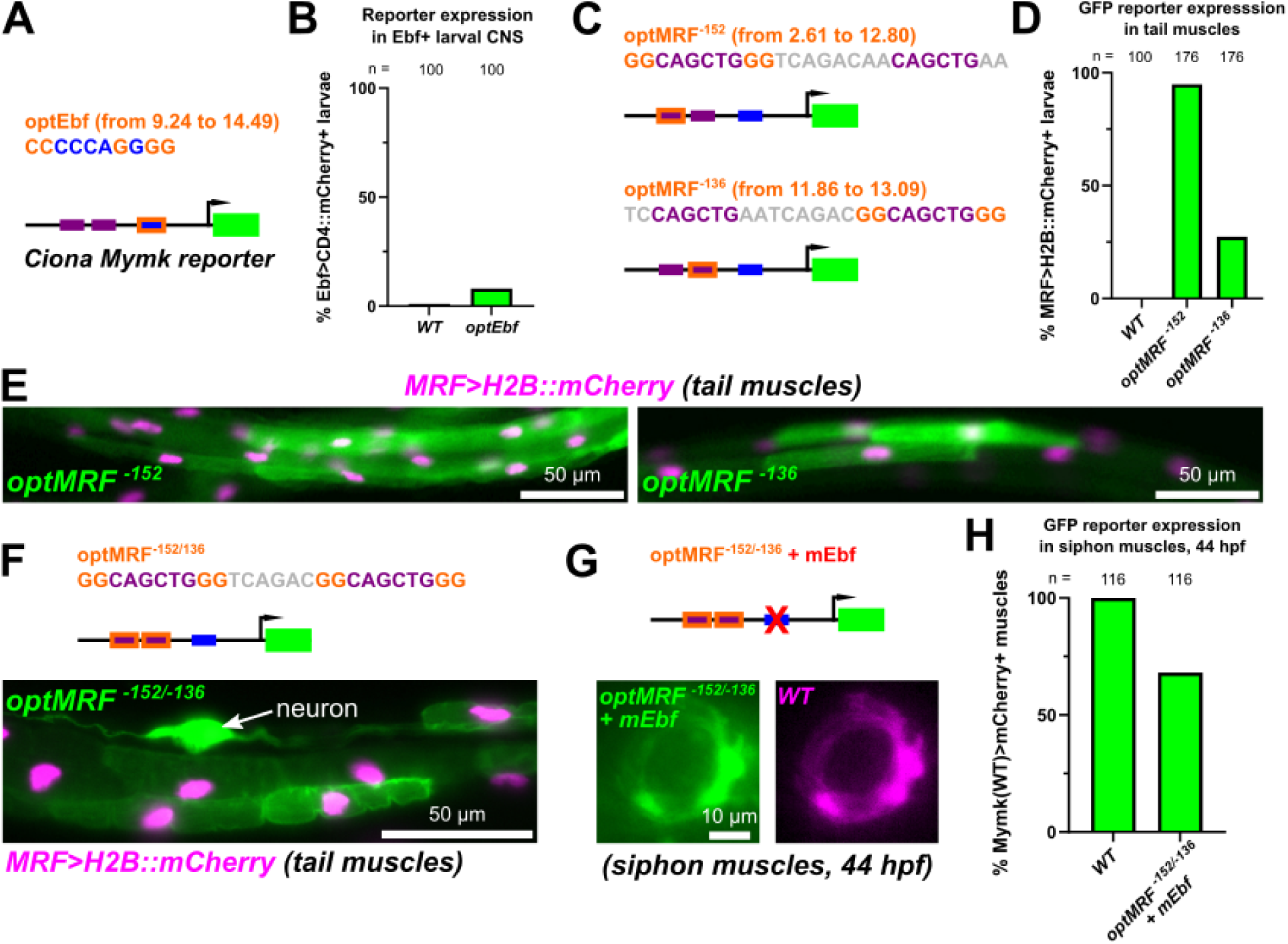
Altered regulatory logic unlocks *Mymk* reporter expression in larval muscles. **(A)** Diagram indicating basepair changes to “optimize” the putative Ebf binding site (optEbf) in the *Ciona Myomaker* promoter by increasing its predicted JASPAR score. (**B**) Optimizing the putative Ebf site (optEbf) resulted in a small but statistically significant (p < 0.0349) effect on activating a *Myomaker* reporter in the absence of MRF in Ebf+ central nervous system (CNS) cells. Larvae fixed at 22 hpf. (**C**) Diagram showing the optimization of either the predicted MRF^-152^ or MRF^-136^ sites. (**D**) Scoring of ectopic reporter expression in larval tail muscles with the optimized putative MRF sites (optMRF), assayed at 17 hpf (p < 0.0001 for both). More frequent ectopic expression was observed with optMRF^-152^ than optMRF^-136^. (**E**) Representative images of larval tail muscles assayed in the previous panel. (**F**) Optimization of both putative MRF^-152^ and MRF^-136^ sites in combination resulted in similar ectopic expression in tail muscles, but also in ectopic expression in neurons in 44% of larvae assayed at 22 hpf. (**G**) Combining optimized MRF^-152^ and MRF^-136^ sites together partially rescues reporter expression even with the putative Ebf binding site disrupted (mEbf). (**H**) Scoring of siphon muscle expression depicted in the previous panel. See text for experimental details, and **Table S1** for statistical test details.

Optimization of the conserved Ebf^-116^ site did not significantly increase *Mymk>GFP* activation in the central nervous system (**Figure 5A,B**). This suggested that either the Ebf site is already “optimal”, or that Ebf binding affinity is not rate-limiting in this context. However, optimization of the conserved, indispensable MRF^-137^ site and/or the non-conserved, dispensable MRF^-152^ site resulted in significant *Mymk>GFP* expression in tail muscles (**Figure 5C-F**). Interestingly, optimization of MRF^-152^ resulted in visible GFP expression in 95% of electroporated larval tails, while optimization of MRF^-136^ resulted in GFP expression in a more modest 27% of tails (**Figure 5D**). Because the increase in average predicted JASPAR score was most pronounced between the wild type MRF^-152^ (JASPAR score 2.61) and its “optimized” counterpart (JASPAR score 12.8, **Figure S3**), this suggested that creating an additional high-scoring MRF binding site is particularly effective for switching a combinatorial MRF-Ebf transcriptional logic to an MRF-alone one. This switch in logic was confirmed when we observed *Mymk* reporter expression in post-metamorphic muscles even when combining optimized MRF sites with a disrupted Ebf site (**Figure 5G,H**). Interestingly, combining both optimized MRF^-152^ and MRF^-136^ sites resulted in ectopic reporter activation in neurons, in addition to tail muscles, in 44% of larvae (**Figure 5F**). This expression might be due to greater affinity for proneural transcription factors that also bind Ebox sequences, such as Neurogenin (Kim et al., 2020). This suggests that the exact sequences flanking each site might also be under purifying selection, minimizing ectopic activation of *Mymk* in tissues where its expression might be detrimental.

## Discussion

In this study, we have investigated the *cis-*regulatory logic of muscle subtype-specific *Mymk* expression in *Ciona*. We have identified two essential transcriptional regulators, MRF and Ebf, that together activate the transcription of *Mymk*, which encodes a transmembrane protein that drives myoblast fusion and muscle multinucleation in tunicates and vertebrates (Zhang et al., 2022). This is in stark contrast to human *MYMK* expression, which only requires the activity of MRFs (Zhang et al., 2020). We have also revealed a potentially conserved repressive input into *Ciona Mymk* transcription, in which direct binding and repression by HES factors might restrict the spatiotemporal window of *Mymk* expression and, consequently, of myoblast fusion. This repression, likely mediated through Delta-Notch signaling, might predate the divergence of tunicates and vertebrates.

We propose that the difference between “MRF+Ebf” and “MRF-alone” logic is responsible for the difference between the pan-skeletal muscle expression of vertebrate *Mymk* and the more selective, post-metamorphic muscle-specific expression in *Ciona* (**Figure 6A**). This in turn might underlie the difference between obligate (vertebrate) versus facultative (tunicate) muscle multinucleation. Although Mymk overexpression is not sufficient to drive the fusion of *Ciona* larval tail muscle cells, *Ciona* Mymk is sufficient to induce human myoblast fusion (Zhang et al., 2022). Our RNAseq results show that this same MRF-Ebf logic is regulating a larger suite of post-metamorphic muscle-specific genes, some of which might encode additional factors required for myoblast fusion.

**Figure 6.**
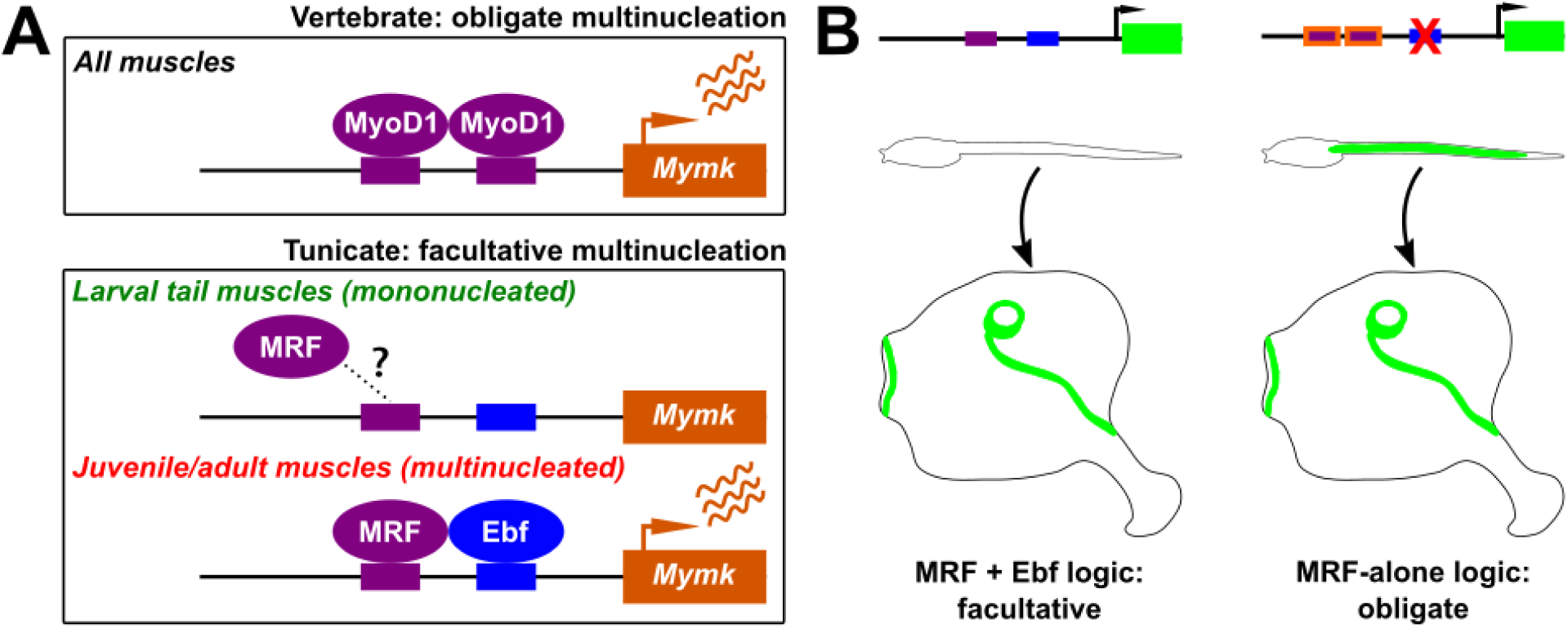
Proposed models for *Mymk* regulation and myoblast fusion in chordates. **(A)** Proposed regulatory models for transcriptional activation of *Mymk* in vertebrates compared to *Ciona* (tunicates). Question mark and dashed lines indicate uncertainty whether MRF can still bind to the *Mymk* promoter in tunicate larval tail muscles, or whether the co-requirement for Ebf acts on other steps independently of MRF binding. (**B**) Summary diagram showing the switch from a combinatorial MRF + Ebf logic to MRF-alone logic for *Ciona Mymk* regulation, obtained in this study through optimization of putative MRF binding sites, with or without disrupting the putative Ebf site.

Although we have largely revealed the basis of *Mymk* regulation in *Ciona*, there are still details that have yet to be elucidated. For instance, what is the mechanism of MRF-Ebf cooperative activity? MRF-Ebf synergy in transcription of muscle subtype-specific genes has been reported in mammals, for instance the activation of the *Atp2a1* gene by MyoD1 and Ebf3 specifically in mouse diaphragm muscles (Jin et al., 2014). Myod1 and Ebf3 and its homologs alone have the ability to activate Atp2a1, but this expression was substantially higher when both MyoD1 and Ebf3 were present. However, there was no evidence of the two transcription factors directly contacting one another to drive cooperative binding. This may be similar to the mechanism of activation of *Mymk* by MRF and Ebf in *Ciona.* Optimization of the predicted Ebf site in the *Mymk* promoter did not significantly increase reporter plasmid expression in Ebf+ larval neurons. This suggests that MRF and Ebf might act on different steps of *Mymk* activation in *Ciona.* In other words, the MRF-Ebf combinatorial logic we have revealed might not be dependent on cooperative binding, as is the case for other examples of cooperativity (Zeitlinger, 2020).

Given the difference between vertebrates (obligate) and tunicate (facultative) activation of *Mymk*, we present two hypotheses for how regulation of *Mymk* might have been controlled in the last common ancestor of tunicates and vertebrates. In the first scenario, the ancestor would have been more like tunicates, in which the combination of MRF and Ebf would have cooperatively activated *Mymk* only in a subset of muscles. In the second scenario, the ancestor would have been more like vertebrates, in which MRF alone would have activated *Mymk* in all muscles.

In the first scenario, the last common ancestor would have had both mononucleated and multinucleated muscles, like we see in most tunicates. It is unclear if the common ancestor had a biphasic life cycle or not, but it is likely they had separate lineages for the trunk and pharyngeal muscles as seen in both vertebrates and tunicates (Razy-Krajka et al., 2014). The ancestor may have had specialized pharyngeal muscles homologous to the siphon muscles of tunicates. One key feature of tunicate siphon muscles is that they are formed by a series of concentric circular myotubes. It is possible that the ancestor had a similar set of circular muscles around the openings of a pharyngeal atrium, and the process of Mymk-driven myoblast fusion might have evolved to allow for the formation of such muscles. After splitting from tunicates, vertebrates would have lost the requirement of Ebf for *Mymk* expression, and MRF would have become the sole activator of *Mymk*, allowing all muscle cells to become multinucleated. This may have been advantageous for their survival, perhaps permitting larger myofibers throughout the body and advanced movement capabilities.

Alternatively, the last common ancestor might have only had multinucleated muscles under the regulation of MRF alone, as in extant vertebrates. Later, vertebrates would have kept this mode of regulation, while tunicates would have recruited Ebf to activate *Mymk* only in post-metamorphic muscles, as an adaptation specifically tied to their biphasic life cycle. As it stands, we do not have enough evidence to conclusively favor one evolutionary scenario over the other. On the vertebrate side, there are no reports of muscle subtype-specific fusion as far as we can tell. On the tunicate side, with the exception of groups that have generally lost the larval phase (e.g. salps and pyrosomes), there are no reports of obligate myoblast fusion. However, we have shown that a switch to pan-muscle expression of *Mymk* is possible through “optimization” of putative MRF binding sites in its promoter, or by creating an additional high-scoring predicted MRF site (**Figure 6B**). Whether this is actually a recapitulation of what happened in evolution or not, we may never know.

## Materials and Methods

### *Ciona* handling, electroporation, fixing, staining, imaging and scoring

*Ciona robusta* (*intestinalis Type A*) specimens were obtained and shipped from San Diego, California, USA (M-REP). The eggs were fertilized, dechorionated, and subjected to electroporation using established methods as detailed in published protocols (Christiaen et al., 2009a, b). The embryos were then raised at a temperature of 20°C. At various stages, including embryos, larvae, and juveniles, the specimens were fixed using MEM-FA solution (composed of 3.7% formaldehyde, 0.1 M MOPS at pH 7.4, 0.5 M NaCl, 1 mM EGTA, 2 mM MgSO4, and 0.1% Triton-X100), followed by rinsing in 1X PBS with 0.4% Triton-X100 and 50 mM NH4Cl to quench autofluorescence, and one final wash in 1X PBS with 0.1% Triton-X100.

Imaging of the specimens was carried out using either a Leica DMI8 or DMIL LED inverted epifluorescence microscope. Scoring was carried out only on mCherry+ individuals as to exclude potentially unelectroporated animals, unless otherwise noted in the figure legends. To carry out CRISPR/Cas9-mediated mutagenesis of *MRF* in the B7.5 lineage we used *Mesp>Cas9* to restrict Cas9 expression to this lineage (Stolfi et al., 2014), together with previously validated *MRF-*targeting sgRNA plasmids *U6>MRF.2* and *U6>MRF.3* (Gandhi et al., 2017). For the negative control, previously published *U6>Control* sgRNA vector was used, which expresses an sgRNA that is predicted to not target any sequence in the *C. robusta* genome (Stolfi et al., 2014). The sgRNAs are expressed *in vivo* from plasmids using the ubiquitous RNA polymerase III-transcribed U6 small RNA promoter (Nishiyama and Fujiwara, 2008). Mutations to disrupt or optimize putative binding sites were all generated through *de novo* synthesis and custom cloning by Twist Bioscience. All GFP or mCherry sequences fused to the N-terminal Unc-76 extranuclear localization tag (Dynes and Ngai, 1998), unless otherwise specified. All plasmid and sgRNA sequences can be found in the **Supplemental Sequences File.** All statistical tests summarized in **Table S1**.

### Ectopic expression of MRF orthologs and *Ciona* Ebf in human *MYOD1*-knockout cells

Human *MYOD1*-knockout myoblasts were generated by CRISPR-Cas9 mediated gene editing and cultured as described previously (Zhang et al., 2020). Retroviral expression vector pMXs-Puro (Cell Biolabs, RTV-012) was used for cloning and the expression of the human *MYOD1*, *Ciona MRF* (Transcript Variant 2), and *Ciona Ebf*. The DNA sequences were verified by Sanger sequencing. For the myogenic rescue experiments, the sgRNA-insensitive version of human MYOD1 open reading frame was used. Retrovirus was produced through transfection of HEK293 cells using FuGENE 6 (Promega, E2692). Two days after transfection, virus medium was collected, filtered and used to infect human myoblasts assisted by polybrene (Sigma-Aldrich, TR-1003-G). When the culture reached 80-90% confluency, cells were induced for myogenic differentiation by switching to myoblast differentiation medium (2% horse serum in DMEM with 1% penicillin/streptomycin). Human myoblasts were differentiated for three days and used for immunostaining and RNA extraction. For immunostaining, the primary antibody for Myosin (Developmental Studies Hybridoma Bank, MF20) and the primary antibody for Myogenin (Developmental Studies Hybridoma Bank, F5D) were used. The qPCR primers for measurements of human *MYMK* and 18S expression are provided in the **Supplemental Sequences File**.

### RNA sequencing and analysis

Total RNA was extracted at 11 hours post-fertilization (Stage 23, late tailbud) from two independent replicates each of electroporated larvae that were transfected either with 50 g *MRF>CD4::GFP* (Negative control) or 50 g *MRF>Ebf transcript variant 1* (Ebf overexpression). Library preparation was at the Georgia Tech Molecular Evolution Core Facility as previously described (Johnson et al., 2023), and sequenced on the Illumina NovaSeq 6000 with an SP PE100bp run. Reads were processed and differential gene expression analysis was performed using DESeq2 in Galaxy as previously described (Johnson et al., 2023). KY21 gene model ID numbers (Satou et al., 2022) were matched to KH gene model ID numbers (Satou et al., 2008) using the Ciona Gene Model Converter application: https://github.com/katarzynampiekarz/ciona_gene_model_converter (Johnson et al., 2023). Our RNAseq analysis was also compared to published microarray analysis of Ebf perturbations in FACS-isolated cardiopharyngeal lineage cells (Razy-Krajka et al., 2014). Volcano plots and comparative transcriptome plots were constructed using R studio and Bioconductor (Huber et al., 2015) with packages EnhancedVolcano (Blighe et al., 2018) and ggplot2 (Wickham, 2016). Raw sequencing reads are archived under NCBI BioProject accession number PRJNA1068599.

## Supporting information

Supplemental Sequences File

## Acknowledgments

The authors are indebted to Dr. Florian Razy-Krajka for thoughtful discussion on Ebf function in specifying post-metamorphic muscle identity in *Ciona*. We thank Lindsey Cohen for technical assistance and Shweta Biliya for help with RNAseq library preparation and sequencing. We thank Brian Hammer, Annalise Paaby, and Will Ratcliff for critical reading and helpful suggestions. This work funded by NIH grant GM143326 to AS, NIH grant GM147209 to PB, and an NSF graduate fellowship to CJJ.

---

**Figure S1.**
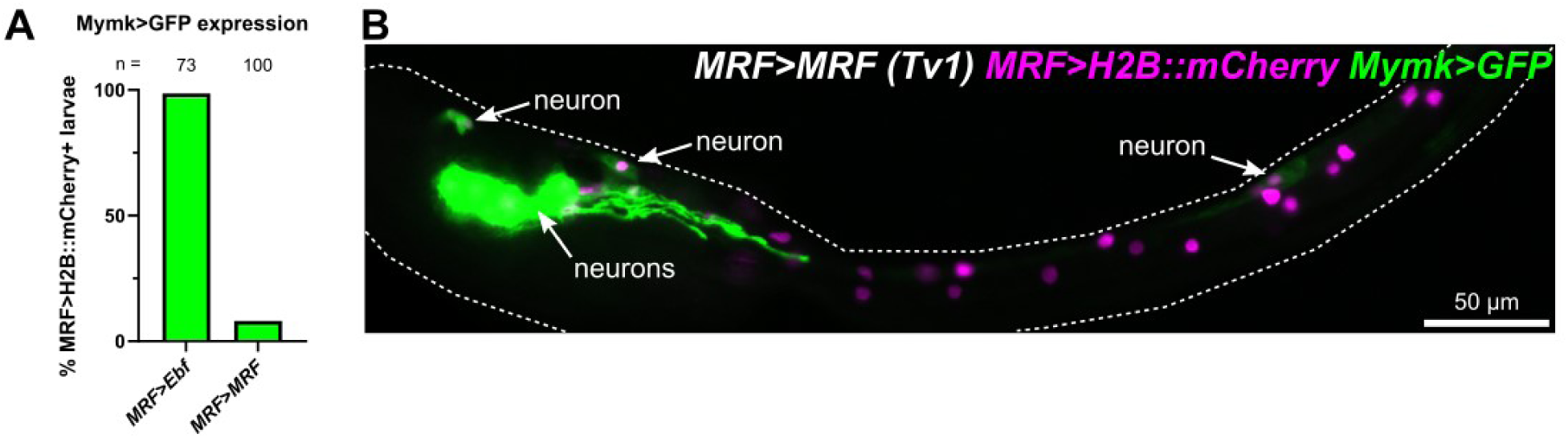
Increased MRF dose in larval tail muscles does not adequately replace Ebf. **(A)** Scoring of larvae at 21 hours post-fertilization (hpf) comparing effect of Ebf or MRF overexpression on ectopic *Mymk>GFP* expression in larval tail muscles (p < 0.0001). (**B**) Most of the effect of *MRF>MRF* on activating ectopic *Mymk>GFP* was limited to the nervous system, likely due to leaky activity of the *MRF* promoter in Ebf+ neurons. See **Table S1** for statistical test details.

**Figure S2.**
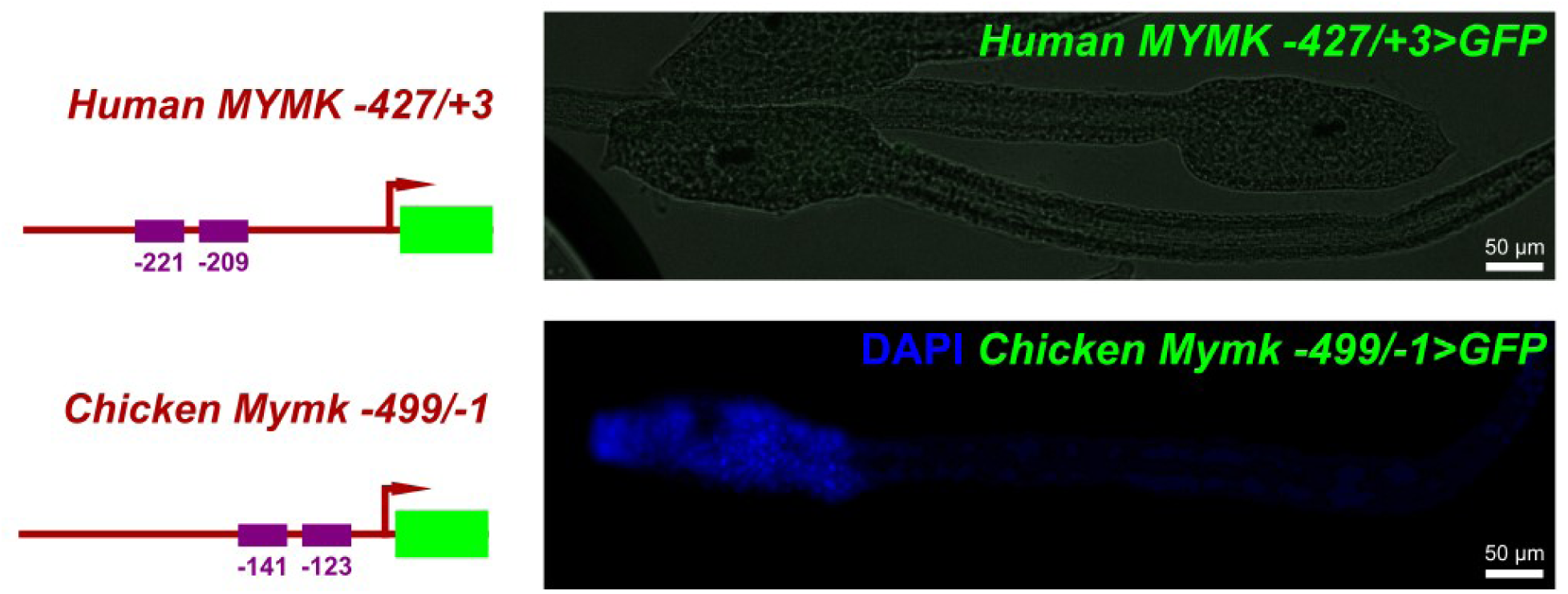
Human *MYMK* and chicken *Mymk* reporter plasmids are not active in *Ciona*. Left: diagrams describing human *MYMK* and chicken *Mymk* reporter plasmids using previously published sequences (Luo et al., 2015; Zhang et al., 2020) and their predicted MRF family member binding sites (purple boxes). Size/spacing of sites is not to exact scale. See **Supplemental Sequences File** for detailed sequences. Right: images of larvae at 17 hours post-fertilization (hpf) electroporated with the reporter plasmids at left, showing no expression in larval tail muscles.

**Figure S3.**
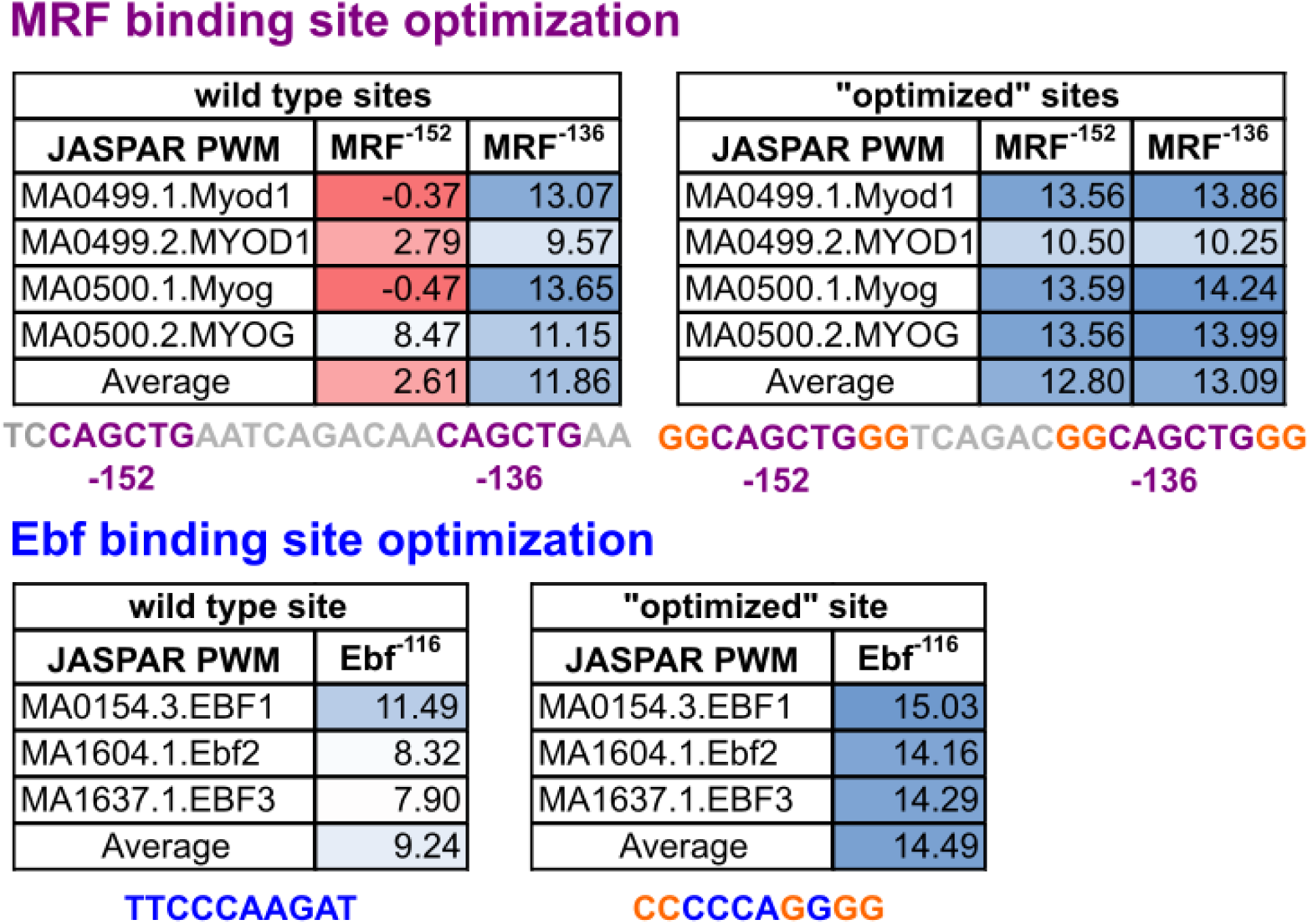
JASPAR scores before and after “optimization” of putative bindings sites. Predicted JASPAR scores for affinity of MRF (top) or Ebf (bottom) to their respective putative binding sites, before and after point mutations to “optimize” them, or rather increase their predicted JASPAR scores. Predictions based on individual human transcription factor PWMs and their averages shown.

**Table S1. Scoring data for *Ciona* experiments and statistical test details**

**Table S2. DESeq2 analysis of differential gene expression of Ebf overexpression in developing larval tail muscles measured by Illumina bulk RNAseq**

**Table S3. Comparison of genes significantly up- or down-regulated in the RNAseq analysis in the current study and the microarray study of Razy-Krajka et al. 2014**

**Table S4. Predicted JASPAR affinity scores for putative MRF and Ebf sites in the *Ciona Mymk* promoter using various human ortholog position weight matrices**

**Supplemental Sequences File. All relevant DNA and protein sequences (reporters, perturbation constructs, primers, etc.) used in this study, including Ciona electroporation mix recipes.**

