## Supplemental Sequences File for "A change in *cis*-regulatory logic underlying obligate versus facultative muscle multinucleation in chordates"

Johnson et al. MRF-Ebf paper

**Relevant coding and non-coding sequences:**

>Ciona Mymk promoter (-508/-1), from Zhang et al. 2022

TGCTCTGGAAAATTTACCAAGGGAAACTCCCTCACGTGGTAACATAGAAATAGTACAAAAAATGCTTAAAATTGTTTTTCACCAAAACTAACGGTAcACATACCGTTGCCTAAACTAGGCAAAACACATATTAGATTGCATTTAAACTGGTAAATGTCCTAATAATTTAAAACTTGCATTAACAATAAATTTAGTTGCCAATAATATAACTTTTAAATTGCGTATTTTAAGTATGTACATAGTTAAGAATATTTATTCGGGCCATAAATGTGTATTTTAAAGTTGTTTATATTTAGACCAAATACTGATATTTTAATAAACGTTTTGGGTCGTGAGAAATATCACATTATTGTGTCCAGCTGAATCAGACAACAGCTGAAGACCACATGCCAATTCCCAAGATAATGGTGCAGCACGCGCAAAACAACCGAGAGTACCGTAGTACATTTGGTATGTCGCCTTTAGAGCATTCATTGCATAGTGTTAAACCGTAGATTAAATCATAGTT

>Ciona MRF promoter (-2604/-1), from Stolfi et al. 2015

gcaagctcctttggggtttggccactcggtcggcaagtaaaatatcatttaagtttttaaatgtcgtttaaatggcagcagccgtaacaggtggactgccacgtgtcgcccggtgtgatcgtgtttccatcccaagggtggtagtgcaggcatgccgacgctcatatgagccattcataataatttggcgcatgtaagaggaaaatgcaccagtaagatggtgccagcacattgtaattaaaatgtggtgctacggagaagtgtggaaatcgttataggaaataaaatcccttattcccgatcgtttttaccaactaacgacgttattttatatttgcgaagttacggttacataattatagtagggtgggggaagatggaacaactttagcacagaatatataaatatcctgatcgagttctaaacaattaacaatggtccatgggttgcgaagatacggttttataattctttgaatgttctttgtttgctaccaaacgggacgagaaaagaaattaaagcatgaaccattttacttcaacctactactacagtatctaagtaatttcttttcgaaacagatttaacttctttgcgatttcaattagcgaatcccagtttgaccgttaaagctatagcttagagtacgtggtgctaaatttcgttgattcgcgcccctgaagcgagcaacttattagatagaaatcccagctggtaacaacatgtgttcatcgccctactgatgtctgtggttgccgaactctctttacgagactgagagccgacgtctcgataaacgaggttgtgtattatgatttatttaggaaataaaaagtaagaaaataaaaatttaattaagtaaaatacaaacgtttacaaaaattatgaatttttttcaccaatttttaattttattcaagtatccgacgacgtttaagaactttaaaatccaaatcctttgctgcttcctagaattatactgtggggtaagatggaaaccgtaaatattctgtgtttacaaccaaataggacgataaaagagaatgaaacgtggacaatttgccctaacacattacacggctaaaaatattaggtttcctcctaaaatagcagcgacgaccataaggttattcgaagatgaaaggccaaagtaacggatgcaagtttttgcaagacaatgctgtaattgttacgtcacagaggaattccacaacttattccagagcagacgagcgtcgcgtttactgtttgctaaaaatggcgactcgaggtgacaattacaattcgcgctgcgaatgacgtaataatgctattcgctgtgatgaaataatagaagtggaaattcgaacaataccaacccacgatgaagcatatgtagcgaatccggaattaaacctctagacttgcttgggaaatgcatggcgtatatgagtgtattacgtgatcgttaaagcgtcttatacagtcgcattagaagctattttaagaaaatccacggcccctcatgcaaatatagctgtgatctaataggttaggttagtttacattaagtttaataatgtcgatcacttattgcatgtttcatgatcttgcaaatatgatgacgttgtatattgaggttgctgctttcagttttatacaatcttccaacaagttaaaactggcgaatttagacaacgattcagtcataaaataaaggccttacgcatctcgagcgaaccagtacgagtgaaaattataacttcaaagtctgcccggcacgtgacttagagtctcccgcgagaccggcgctccggagttctggaatgcagaagaagaagaagcattgtcacctaccgtgacgtcataaacgagtgtcttaaatagcgcctgcatcgagccatgcattctataacatcgctagccagccacgcatggtaaagttatcaagcgatagccagtcaatgatatatagtagccaatgtctgttataaccaggcatggtcacatagcgtgttttaatacaagtagatttacatggtatatagttagcggcgaactaatgttttaaactgcagcaaattgcagacattaggtcttgcaaaagtagcctacgagtattacattagccattcggtgatcaagttattttacacagaattcatacttaatgtttagtttgggagttcgtaataatatgttaattgcaattaaactgcaagccttatcgttgcacgaagtatattatataccagactttactacatatagaatcagaaagaatacagttacgaatcgccgacgaaacaagcttctaatattaaatcgtgaaacattataaaagtgaatggattaaacttatcacgcttaaattatatcaagtcttcagggtatatacttttcgccggttctaaattatttttcgcaagacttgttttttatacaaaatcttaacctaaacgaatgtgcagttataggtttaatattatgtgttacttgcaattacagtgcaaggagaaccgttgcaaaattacaaccataagttgcaagatattaagattatgtattgacctcatacattttgtatttcagaaatctagccggtagtttgacatatttatacg

>Ciona MRF promoter (-2604/-6 version)

gcaagctcctttggggtttggccactcggtcggcaagtaaaatatcatttaagtttttaaatgtcgtttaaatggcagcagccgtaacaggtggactgccacgtgtcgcccggtgtgatcgtgtttccatcccaagggtggtagtgcaggcatgccgacgctcatatgagccattcataataatttggcgcatgtaagaggaaaatgcaccagtaagatggtgccagcacattgtaattaaaatgtggtgctacggagaagtgtggaaatcgttataggaaataaaatcccttattcccgatcgtttttaccaactaacgacgttattttatatttgcgaagttacggttacataattatagtagggtgggggaagatggaacaactttagcacagaatatataaatatcctgatcgagttctaaacaattaacaatggtccatgggttgcgaagatacggttttataattctttgaatgttctttgtttgctaccaaacgggacgagaaaagaaattaaagcatgaaccattttacttcaacctactactacagtatctaagtaatttcttttcgaaacagatttaacttctttgcgatttcaattagcgaatcccagtttgaccgttaaagctatagcttagagtacgtggtgctaaatttcgttgattcgcgcccctgaagcgagcaacttattagatagaaatcccagctggtaacaacatgtgttcatcgccctactgatgtctgtggttgccgaactctctttacgagactgagagccgacgtctcgataaacgaggttgtgtattatgatttatttaggaaataaaaagtaagaaaataaaaatttaattaagtaaaatacaaacgtttacaaaaattatgaatttttttcaccaatttttaattttattcaagtatccgacgacgtttaagaactttaaaatccaaatcctttgctgcttcctagaattatactgtggggtaagatggaaaccgtaaatattctgtgtttacaaccaaataggacgataaaagagaatgaaacgtggacaatttgccctaacacattacacggctaaaaatattaggtttcctcctaaaatagcagcgacgaccataaggttattcgaagatgaaaggccaaagtaacggatgcaagtttttgcaagacaatgctgtaattgttacgtcacagaggaattccacaacttattccagagcagacgagcgtcgcgtttactgtttgctaaaaatggcgactcgaggtgacaattacaattcgcgctgcgaatgacgtaataatgctattcgctgtgatgaaataatagaagtggaaattcgaacaataccaacccacgatgaagcatatgtagcgaatccggaattaaacctctagacttgcttgggaaatgcatggcgtatatgagtgtattacgtgatcgttaaagcgtcttatacagtcgcattagaagctattttaagaaaatccacggcccctcatgcaaatatagctgtgatctaataggttaggttagtttacattaagtttaataatgtcgatcacttattgcatgtttcatgatcttgcaaatatgatgacgttgtatattgaggttgctgctttcagttttatacaatcttccaacaagttaaaactggcgaatttagacaacgattcagtcataaaataaaggccttacgcatctcgagcgaaccagtacgagtgaaaattataacttcaaagtctgcccggcacgtgacttagagtctcccgcgagaccggcgctccggagttctggaatgcagaagaagaagaagcattgtcacctaccgtgacgtcataaacgagtgtcttaaatagcgcctgcatcgagccatgcattctataacatcgctagccagccacgcatggtaaagttatcaagcgatagccagtcaatgatatatagtagccaatgtctgttataaccaggcatggtcacatagcgtgttttaatacaagtagatttacatggtatatagttagcggcgaactaatgttttaaactgcagcaaattgcagacattaggtcttgcaaaagtagcctacgagtattacattagccattcggtgatcaagttattttacacagaattcatacttaatgtttagtttgggagttcgtaataatatgttaattgcaattaaactgcaagccttatcgttgcacgaagtatattatataccagactttactacatatagaatcagaaagaatacagttacgaatcgccgacgaaacaagcttctaatattaaatcgtgaaacattataaaagtgaatggattaaacttatcacgcttaaattatatcaagtcttcagggtatatacttttcgccggttctaaattatttttcgcaagacttgttttttatacaaaatcttaacctaaacgaatgtgcagttataggtttaatattatgtgttacttgcaattacagtgcaaggagaaccgttgcaaaattacaaccataagttgcaagatattaagattatgtattgacctcatacattttgtatttcagaaatctagccggtagtttgacatattt

>Ciona Mesp promoter (-1914/-1) from Davidson et al. 2005

cggttcaacgtgacgtcccatgccgatatcgtaaccatccggaacctctgatgctttttcaatatcatcttttttgaaatccttcatttccgtttcatcgtcatttttgaaagccgggttctcatactcgtttttggtgctgtaaccgttttctgacatttttatctcatccaagtgcgaaccacttcaaagatggatagataccagattttattacaaacaattagcaattcacgaaagttaaaaaaacgacataaaactataaaataaaattatacttattaaatatgaagaaaaatatgcatttttaatcattctattgcaacaaatcggtatttttcgtattctcttatacgaagattgcatacaagcttaacgtttcatctgtttccattggaataaaatagaaaacgtcccaccgcctgccctaactttttataagcttcacaaattatcaaagtttaccgttgaactcctgataaatgattattcagctattttaaacccaacttcgtttataagtgatgccgtctccttcttcgtatgaccccgctctactaaatacatgtttgcacactgatcctaatgaagaccgcacctgaacctttacatagccgatgcattcacatataatgttctttgtgcaaccgaagtcgtatttgagcaaaccaaatatccaacttatataagtagacctttaaaaatggacggtgaacaaaactttgcgaagtcatcataattgaaggtttatttagctctcctattgtagggaaagatcctatggatgctccaaaacgataaacccgacttaccgaaaacgcggcggtcgttagaccatgcttattgataaaccacatacatacaaactttagcgcaaactcggctatatctttacaacattagacaaattctctttcatgtattttcatgggattataaagtgtctattttaggtaacaagactttaataaaaatatcagtgtcccatattgttggcggctttgtttgcgagtgtctctgtattgacaacgtttataattaacttggaagtaatgagtatacagcagcagacacagaagtttgtcgggggtatcctatgattgtacatcatgtgagcaaatatacttcaactgctagtgaataatatttgatatacgcacaagtcgtacaaatacacgaccacaaataagtcatacttgtcgaagtttgtattcccctgttttcgatcaatcttatcagccacaaaaaatggaaaaattccaaaaacgtagacacccacaaagtaacatacatggtaactcgtaagctaatacagagcacccgtgtgaaaacgactgtcgttttccggccacgcgagaataaagtaagttacattgattcattcactgacagtttcccaccaaaactgaagtattaatcaattatgaatcggcagtgctttcagtgggaagatttcggaaaatgtgggtctaatgtcacagtaaactttgtgtttgttttaaaaacaatcatcgttaaatcaacatcgcaattctgtcaatcactccacaataaactatatggctaatggaaacaggctgtttgtttgtatatcataaaatacttgcgtcgtattatctgaccaaacaaaagcgttacaggaacggtccagcttcaaaatttgctgatgcatcagagtttacatttgaaatgtgattaattacgaaaatccagcgaatagaattgtcacaacaagtcattagcgacggatatttcgcctttgaaacttaaaggcgataatgactttgcccgtttcatgcggcgataaacgaactaattagacacctcctacagatataatggtaattcagaatcgtgtggttatgtaattcgcaaaaacattttaacaaaacaggttgatttgaaacttgtatt

>nls::Cas9::nls from Stolfi et al. 2014 START STOP

ATGGCTAGCCCCAAAAAGAAGAGGAAAGTGGACAAGAAGTATTCTATCGGACTGGACATCGGGACTAATAGCGTCGGGTGGGCCGTGATCACTGACGAGTACAAGGTGCCCTCTAAGAAGTTCAAGGTGCTCGGGAACACCGACCGGCATTCCATCAAGAAAAATCTGATCGGAGCTCTCCTCTTTGATTCAGGGGAGACCGCTGAAGCAACCCGCCTCAAGCGGACTGCTAGACGGCGGTACACCAGGAGGAAGAACCGGATTTGTTACCTTCAAGAGATATTCTCCAACGAAATGGCAAAGGTCGACGACAGCTTCTTCCATAGGCTGGAAGAATCATTCCTCGTGGAAGAGGATAAGAAGCATGAACGGCATCCCATCTTCGGTAATATCGTCGACGAGGTGGCCTATCACGAGAAATACCCAACCATCTACCATCTTCGCAAAAAGCTGGTGGACTCAACCGACAAGGCAGACCTCCGGCTTATCTACCTGGCCCTGGCCCACATGATCAAGTTCAGAGGCCACTTCCTGATCGAGGGCGACCTCAATCCTGACAATAGCGATGTGGATAAACTGTTCATCCAGCTGGTGCAGACTTACAACCAGCTCTTTGAAGAGAACCCCATCAATGCAAGCGGAGTCGATGCCAAGGCCATTCTGTCAGCCCGGCTGTCAAAGAGCCGCGGACTTGAGAATCTTATCGCTCAGCTGCCGGGTGAAAAGAAAAATGGACTGTTCGGGAACCTGATTGCTCTTTCACTTGGGCTGACTCCCAATTTCAAGTCTAATTTCGACCTGGCAGAGGATGCCAAGCTGCAACTGTCCAAGGACACCTATGATGACGATCTCGACAACCTCCTGGCCCAGATCGGTGACCAATACGCCGACCTTTTCCTTGCTGCTAAGAATCTTTCTGACGCCATCCTGCTGTCTGACATTCTCCGCGTGAACACTGAAATCACCAAGGCCCCTCTTTCAGCTTCAATGATTAAGCGGTATGATGAGCACCACCAGGACCTGACCCTGCTTAAGGCACTCGTCCGGCAGCAGCTTCCGGAGAAGTACAAGGAAATCTTCTTTGACCAGTCAAAGAATGGATACGCCGGCTACATCGACGGAGGTGCCTCCCAAGAGGAATTTTATAAGTTTATCAAACCTATCCTTGAGAAGATGGACGGCACCGAAGAGCTCCTCGTGAAACTGAATCGGGAGGATCTGCTGCGGAAGCAGCGCACTTTCGACAATGGGAGCATTCCCCACCAGATCCATCTTGGGGAGCTTCACGCCATCCTTCGGCGCCAAGAGGACTTCTACCCCTTTCTTAAGGACAACAGGGAGAAGATTGAGAAAATTCTCACTTTCCGCATCCCCTACTACGTGGGACCCCTCGCCAGAGGAAATAGCCGGTTTGCTTGGATGACCAGAAAGTCAGAAGAAACTATCACTCCCTGGAACTTCGAAGAGGTGGTGGACAAGGGAGCCAGCGCTCAGTCATTCATCGAACGGATGACTAACTTCGATAAGAACCTCCCCAATGAGAAGGTCCTGCCGAAACATTCCCTGCTCTACGAGTACTTTACCGTGTACAACGAGCTGACCAAGGTGAAATATGTCACCGAAGGGATGAGGAAGCCCGCATTCCTGTCAGGCGAACAAAAGAAGGCAATTGTGGACCTTCTGTTCAAGACCAATAGAAAGGTGACCGTGAAGCAGCTGAAGGAGGACTATTTCAAGAAAATTGAATGCTTCGACTCTGTGGAGATTAGCGGGGTCGAAGATCGGTTCAACGCAAGCCTGGGTACCTACCATGATCTGCTTAAGATCATCAAGGACAAGGATTTTCTGGACAATGAGGAGAACGAGGACATCCTTGAGGACATTGTCCTGACTCTCACTCTGTTCGAGGACCGGGAAATGATCGAGGAGAGGCTTAAGACCTACGCCCATCTGTTCGACGATAAAGTGATGAAGCAACTTAAACGGAGAAGATATACCGGATGGGGACGCCTTAGCCGCAAACTCATCAACGGAATCCGGGACAAACAGAGCGGAAAGACCATTCTTGATTTCCTTAAGAGCGACGGATTCGCTAATCGCAACTTCATGCAACTTATCCATGATGATTCCCTGACCTTTAAGGAGGACATCCAGAAGGCCCAAGTGTCTGGACAAGGTGACTCACTGCACGAGCATATCGCAAATCTGGCTGGTTCACCCGCTATTAAGAAGGGTATTCTCCAGACCGTGAAAGTCGTGGACGAGCTGGTCAAGGTGATGGGTCGCCATAAACCAGAGAACATTGTCATCGAGATGGCCAGGGAAAACCAGACTACCCAGAAGGGACAGAAGAACAGCAGGGAGCGGATGAAAAGAATTGAGGAAGGGATTAAGGAGCTCGGGTCACAGATCCTTAAAGAGCACCCGGTGGAAAACACCCAGCTTCAGAATGAGAAGCTCTATCTGTACTACCTTCAAAATGGACGCGATATGTATGTGGACCAAGAGCTTGATATCAACAGGCTCTCAGACTACGACGTGGACCACATCGTCCCTCAGAGCTTCCTCAAAGACGACTCAATTGACAATAAGGTGCTGACTCGCTCAGACAAGAACCGGGGAAAGTCAGATAACGTGCCCTCAGAGGAAGTCGTGAAAAAGATGAAGAACTATTGGCGCCAGCTTCTGAACGCAAAGCTGATCACTCAGCGGAAGTTCGACAATCTCACTAAGGCTGAGAGGGGCGGACTGAGCGAACTGGACAAAGCAGGATTCATTAAACGGCAACTTGTGGAGACTCGGCAGATTACTAAACATGTCGCCCAAATCCTTGACTCACGCATGAATACCAAGTACGACGAAAACGACAAACTTATCCGCGAGGTGAAGGTGATTACCCTGAAGTCCAAGCTGGTCAGCGATTTCAGAAAGGACTTTCAATTCTACAAAGTGCGGGAGATCAATAACTATCATCATGCTCATGACGCATATCTGAATGCCGTGGTGGGAACCGCCCTGATCAAGAAGTACCCAAAGCTGGAAAGCGAGTTCGTGTACGGAGACTACAAGGTCTACGACGTGCGCAAGATGATTGCCAAATCTGAGCAGGAGATCGGAAAGGCCACCGCAAAGTACTTCTTCTACAGCAACATCATGAATTTCTTCAAGACCGAAATCACCCTTGCAAACGGTGAGATCCGGAAGAGGCCGCTCATCGAGACTAATGGGGAGACTGGCGAAATCGTGTGGGACAAGGGCAGAGATTTCGCTACCGTGCGCAAAGTGCTTTCTATGCCTCAAGTGAACATCGTGAAGAAAACCGAGGTGCAAACCGGAGGCTTTTCTAAGGAATCAATCCTCCCCAAGCGCAACTCCGACAAGCTCATTGCAAGGAAGAAGGATTGGGACCCTAAGAAGTACGGCGGATTCGATTCACCAACTGTGGCTTATTCTGTCCTGGTCGTGGCTAAGGTGGAAAAAGGAAAGTCTAAGAAGCTCAAGAGCGTGAAGGAACTGCTGGGTATCACCATTATGGAGCGCAGCTCCTTCGAGAAGAACCCAATTGACTTTCTCGAAGCCAAAGGTTACAAGGAAGTCAAGAAGGACCTTATCATCAAGCTCCCAAAGTATAGCCTGTTCGAACTGGAGAATGGGCGGAAGCGGATGCTCGCCTCCGCTGGCGAACTTCAGAAGGGTAATGAGCTGGCTCTCCCCTCCAAGTACGTGAATTTCCTCTACCTTGCAAGCCATTACGAGAAGCTGAAGGGGAGCCCCGAGGACAACGAGCAAAAGCAACTGTTTGTGGAGCAGCATAAGCATTATCTGGACGAGATCATTGAGCAGATTTCCGAGTTTTCTAAACGCGTCATTCTCGCTGATGCCAACCTCGATAAAGTCCTTAGCGCATACAATAAGCACAGAGACAAACCAATTCGGGAGCAGGCTGAGAATATCATCCACCTGTTCACCCTCACCAATCTTGGTGCCCCTGCCGCATTCAAGTACTTCGACACCACCATCGACCGGAAACGCTATACCTCCACCAAAGAAGTGCTGGACGCCACCCTCATCCACCAGAGCATCACCGGACTTTACGAAACTCGGATTGACCTCTCACAGCTCGGAGGGGATGAGGGAGCTCCCAAGAAAAAGCGCAAGGTAGGTTAATGA

>Ciona U6 promoter from Nishiyama and Fujiwara 2008

tggcgggtgtattaaaccactaaacaaacaattgccccaagctctcttcacaattataaacactacaaatgtttggacaagagattagcgtggctgtgacgagtaatctcaaaggcttggtgtaattgatattttataagaagcagattaaacttcaatacagttaacacctcatttacaaaaaattggctgccaaaatcgctaattaacacatatttaaaacaatttctacagatatacacagtatagtatgattactaactgcataataaacaaacatatccaacagacactcactaatctgccataacaagcttcaaaaacttaaactcgaaattttagtgaatctttttttttaatgaagattttatttaaaaagttaaaaatattacagttcaggtataggtttacacctaatctttaataatccgaactaaattttaactatttagaaactttttcaaccaaagtttaaaaaaatagattcttcgcacgctaaaactatcatttacacaaaaaaatgcaacaaaatgcagaaaaaaattacattagagtttaggttagttacctgctaatcattataaactaacttccggcataatattcatctaaaattagcaataatcacgttttacgctaaaatttgtgtaaaactaaacttcgtcctttgtcaaggagaaaatttgactcaaaagctgcgcgcgcaggggagatccccaagcgagtgtttgttacatcataatcatgtggaaaaatcccctaataagtaaaaatacatattttttaattttgggggcaaataaaccgctttttatgtctaaaaacgccaaaaatggatcgcgcgagcccaaaaacgcacaaataacgtacagacagtgtctctgcgtacacagacggtatttcccctttaaattgagaactagacttaagcacgcttataagtctggaaggcatccgatggtatagat

>Ciona Ebf promoter (-2631/+15+STOP) START from Stolfi and Levine 2011

attcttccgggaaataagagcggcagcgactttattcgaatttcaaatttccatgatattcgataccccacactccaccctataacggtgacctggtttaggaaatatgtgacgttgcttcacgacaattttagcagctatataagttaccattgcaaaaagatcagtctttttatttatcacaatttttaccagatttttaaatattttttagaaataacttaaattttgatcaaactgggttaaaattgcaaattttacatttttgtgattaatattaatctttttagtcgccatggcttaaaaatgtcatttgtttagtccgctgtattcaattttaccgtttgataaatcaaacggttaagtttcgcataaaaccgatttctcatatggatattactccgcgcgagttctaacaacgcgttctcttcgctggccaagatcgatactccgtgacgtcacaatcacatgcgcgcgcgtctgaaatatggtcgacagagttgcaacacgcaacccttttgtttgggttaatttccctctttgttttttaagtccgatcgcgtcgccacatctgttcgacgaatcttcgttctgtcttagcaactttgcggctctgtccattgtgaagtcgtaaacgaggcattgtcgtcactgctgtctcgcctacgtcacaaagcagtgcacagtgacgtcacggagacgaggtcgcgtgccttcgagttggaaaattcaaaccattgcactttttgccgaatgttaatttttacgacgggagaagttaggaaaagcataaaataacaaataaataaccaataaaaaacgtaaaaacaaacaagagaacggtattcaaaattagaatttcgaaaaaaatgttaaaaaataatacttagagtcgctgaacaatttaagccgacaaaggccacaaaactgcctaaaatttataaaaaaaaatgtaaaattattttgttttttttaaactacagttatcacctttaaaacaaacaaattagcaaacgttgtaattacttcacaactttcttgcgacgctaaaaggcggcgaattttattgctattgtgacgtcacaagcgctctcgtcacgcccggatacgattagaacaacgaaggattgtttgtttttaattatttctctgtttaatcatttgatttagcgcggcacaaattttgttttatataaaatgtatctatttatccattttatttctgtgcgttgttgtactattttttgaaaaatgtttgttaacctttagaaaatcgcgaaaccaacgaaattatttctaaaggctgtacaattctttttcgttaggttacgtgtttaagtataggaccaagttttaaatggcggacagagtttcgtattttgatatcttgaattattttgaaatcttgaaaaaaaataattgtacgctttacatagaataactaaccaaaatatctgaagcaaaaattacgtacaattttttaaaatgtaatttactttctagcttttaatttttgtgcttttctcaatatgtgtgccatattttaaaaacgtaaatttgcctgttgttaagcggaggaaaaagtaatctgcgtgaatgcgaaaaacacgattttggaatcagcgccggcaatggtgtttgtaaataggggtggcatacgcgtttcgtagcgaaagagagaatgaggcgaaagtcgacagatgcacgctccgatttatgagacaggaaccagtcgcgagggcacggaggaaaaagaaccttactccaagacatgcgccgccttttttctttctgtcctagtcaggaatactagagtatagaaggccacgcgtcgtggagtttaaaaccagcaagcgagtgtctcaacggacacatcaatacagagacttctctcagtggacaactcggacgattcgccactaacttggtggattcgtcccggacgacccatgggccgggtcccagcgcgctagttggccaccatacagtgtagaatcagctagatcgtctctgcggatttcgcaaatagattgagttggagataggttcccgaccgggatttcgactaatttgcaatgttagttattaatcaaggtgacagtcaggagttaaagttaatttaccttttgaaagggcaaaaagattttcgaagtaaattgattccgttaattgtaactcttaaacgcaaaccgataaacgacgccattttgcttttcattgagaaactaaccatttaggcattctataattaaaattaaatgtttttaaaatctgtaactatctgtaataaactagaattatttatttcagtttaaatttttatttaaaacaataattttattatttcctttattcatcctattgatattggtatagagaaaacgatgtttttatttccaaaaaattttgtttcataaaagtcattttgttcaatttaaaaaaataccttattctaaaaaatgcccaaaacaagttctttatttcttaaaaactatcctaaataaaaaaaaactccacgttttaataaaacctataaaattaaaaactataaaaagatgacttatttttttaccctaacgtgatttttcaccagataccttaagtgttattttatttgtaagtaatatccaaATGGCAACAATCGCGTAA

>Ciona Ebf START STOP

ATGGCAACAATGGCGGGACCTCAGTTATCGGGGCCAGCAGTGAGAGGATGGATGCAAACAACGTTAGTGGAACCGATGCCAAACGGCAATGTTGGTCTACATCGGGCACATTTCGAAAAGCAGCCGCCAAACAACCTTAGAAAAAGCAACTTTTTCCACTTCGTGCTTGCCCTGTACGACAGACAAGGGCAGCCGGTGGAAATTGAGCGAAGTGCTTTCGTAGGATTTGTGGAAAACGAAACCGAGATCGCCGGCGAAAAAACAAACAACGGGATCCAGTACAGACTCCAATTGCTTTACCATAGCGGCGTACGGACAGAACAAGATGTATTCGTCCGTCTTATCGACTCTGCAACCAAACAGTCAATTACGTACGAAGGACAGGATAAAAACCCAGAAATGCGACGGGTTTTGCTCACTCATGAGATCATGTGCAGCCGCTGCTGTGACAAGAAGAGTTGCGGGAACAGAAACGAAACTCCATCCGACCCAGTGGTGATTGACAGATACTTCTTAAAATTCTTTCTCAAGTGCAACCAAAACTGTTTAAAGAATGCTGGAAATCCAAGAGACATGCGAAGATTTCAGGTTGTGGTCTCAACAACTGTGCATGTTGACGGTCATGTTTTAGCTGTGTCTGACAACATGTTCGTGCACAACAACTCGAAACATGGACGTCGAGCGAGAAGGGTCGACCCATCAGAAGCTTCGCCAACTATCAAAGCAATCAACCCAGCTGAGGGTTGGACAACAGGTGGAGCAACAGTTGTTATAGTTGGAGAAAATTTCTTTGACGGGCTTCAAGTTGTGTTTGGTTCAATGGTTGTATGGAGTGAGTTGATCACCCAACACGCAATCAGAGTACAGACTCCACCGCGCCATCTGCCGGGAGTGGTAGAAGTGACTCTGTCTTACAAAAATAAACAATTCTGCAGTGGTGCTCCAGGACGCTTTGTATACACAGCGCTCAACGAACCGACACTTGATTATGGATTCCAGCGATTACTGAAGACTGTGCCAAGACACCCAGGCGACCCTGAGAGGCTACCAAAGGAGATTATTCTAAAACGAGCTGCTGACGTCATGGAAGCGGTGATCAGTCGGCAGTACGCCCCGCCAAGCCAAATGCCCCCGTCAGCCGGGATCACGCCTCCGGCGCCCCATCTAGCCGCGGCACCGTGCGCACCACCTGGCAGTTTCGTTCCGCAATCTGCTAGCGCTGCCATGGCCGTGGCAATGAACGGTTACGCTGCTGCAGCGGTTTCTTCACAATTTGGCGGAACACCCGATCGGTTCGACACCGGAAGTGATTCAGGTTACTCACGAGGTAACAGTGTTTCTCCCCGCAACGGATATTCCCCCCAGACGACACCGCATAGCTTGAACAGTGGCTCAATTGGTAGCATGGTGGGGCTCACAACTGTAGGCGCAGTTCCGGCCCCAGCCCCCTACCACTGCGCACCATCCTTTAACAGTTATTCCAGCGCTTCAGGTCCCACATTCACGAACATGAACAACACGTCTCCTGGCCTTTTTTCTGGATCTGGGATAATTCCACCTTCGCCCCACAACGGTATGAACCCACTCCCGTCTTCGGGGACCACACCGGGAATTTTTAGTTTTTCACCTGCAAACATGATATCAGCCGCAAAGCAGAAGAGCGCATTTGCCCCTGTACATCGTCCACACAACTCGCCCAGTCCTCTAGCGCCATCTAACGGAAATATCGCACTAAACGGCTATAGCTAG

>Human EBF3 (codon optimized for C. elegans) START STOP

ATGTTCGGAATCCAAGAAAACATACCAAGAGGCGGTACTACTATGAAAGAAGAACCACTTGGATCAGGAATGAATCCAGTTAGATCATGGATGCATACTGCTGGAGTCGTTGATGCGAATACAGCTGCGCAATCTGGAGTTGGACTTGCTAGAGCACATTTTGAGAAACAACCACCAAGTAATTTACGAAAGTCAAACTTCTTTCATTTTGTCTTGGCCCTTTATGACAGACAAGGTCAACCTGTAGAAATCGAGAGAACAGCGTTCGTTGATTTCGTTGAAAAGGAAAAGGAACCTAATAATGAGAAGACTAATAATGGTATCCATTACAAGCTTCAATTGTTATATAGTAATGGGGTTAGGACCGAGCAGGACCTTTACGTCAGACTGATTGACAGCATGACTAAGCAAGCAATTGTTTATGAAGGGCAAGATAAGAATCCTGAAATGTGTAGGGTTTTGCTTACGCATGAAATTATGTGTTCGCGTTGTTGCGATAAGAAGTCTTGCGGAAACCGAAATGAGACCCCTAGTGATCCAGTCATTATAGATAGGTTCTTCCTGAAATTCTTTCTTAAATGTAACCAAAATTGCCTTAAGAACGCCGGAAATCCACGTGACATGCGCCGGTTTCAAGTAGTAGTCTCTACCACCGTAAATGTTGATGGACATGTATTGGCTGTTTCTGATAATATGTTCGTTCATAATAACTCAAAGCATGGACGACGAGCACGTCGTTTGGATCCAAGCGAGGGAACAGCTCCATCATACCTTGAGAACGCAACACCATGTATTAAAGCTATTTCTCCGTCCGAGGGATGGACGACTGGCGGCGCAACAGTGATTATTATCGGAGATAATTTCTTCGATGGCCTTCAGGTGGTTTTCGGTACGATGCTTGTTTGGAGTGAATTGATCACACCTCACGCGATTCGGGTTCAAACGCCACCTCGTCATATCCCGGGAGTTGTTGAGGTAACGCTTTCTTATAAGTCGAAACAATTTTGTAAGGGAGCACCGGGCAGGTTCGTATATACTGCATTAAACGAGCCGACAATTGACTATGGATTCCAACGTCTGCAAAAGGTTATACCGCGTCACCCAGGAGACCCAGAGCGCTTGCCAAAAGAAGTTCTTTTGAAAAGGGCCGCTGATTTGGTAGAGGCATTGTATGGTATGCCACATAATAATCAAGAAATTATATTAAAGAGAGCCGCTGATATTGCAGAGGCTCTCTATTCTGTCCCAAGGAACCATAATCAAATACCAACACTTGGAAATAATCCAGCCCATACCGGAATGATGGGAGTTAATAGCTTTTCGTCACAACTGGCTGTGAATGTCTCGGAAACTTCCCAGGCTAATGATCAGGTGGGGTATTCACGGAACACCTCTTCGGTCTCACCAAGAGGATATGTTCCGTCATCCACGCCACAACAATCAAACTATAATACTGTTTCAACCAGTATGAACGGGTACGGGAGCGGAGCGATGGCAAGCCTTGGAGTACCGGGATCACCGGGTTTCCTGAACGGTTCAAGTGCCAATTCACCATATGGAATCGTTCCCTCAAGTCCTACAATGGCTGCTTCATCTGTAACGCTTCCATCCAATTGCTCATCCACTCATGGAATCTTTTCGTTTTCTCCGGCTAACGTAATTTCAGCGGTCAAGCAAAAGTCAGCATTTGCACCAGTTGTTAGACCACAGGCTTCCCCACCACCATCATGTACGTCTGCGAATGGAAACGGTTTGCAGGGAAGCCTTCTCGGAGCGGAAGATGTGGCGGCTGAGAAAACGAATTGGCCTTTCTGCGAGGTTGGTGGGATTTTCCATTTCGACGAGCTTATGCTAAAGAAGGGCACAGGGAAATTGTGCTTAGGTTGGTAA

>Ciona MRF (transcript variant 1, with sgRNA mismatches) START STOP

ATGACGTGTATCTCTCTAGAGGAGCTCGACCTCTCTTCAATATTCTCCAACAGCAGCAGCTATTTCACCAGTTACGCCACAAACCCCATTATGACGTCACAAAAACGGACGCCTTCACGACTGAAGCGAGCAAGCAGCGACGTTTTGTTGTCAGATACAGCGGTTTCGCCGGTAAGTCGAGAAGTTTCGGGGGTTTTATCGGAACTGGACGAACTGAAGAGATGtGTcGAGGGAAATTATATCGGAATCGATGCGGAAAAGAGCGATATCTCGATACTTGAGGAACTTAGTAATATGGCGTCCGGTTGTAGCGATTCAGACGTCGCTTACTCGTCCCCGGATTCGAGGTACGGGTCAACTGGTAACCTTACCAGCAAGACAAGTTTCGGATCGGGAATGGGTTTCCGTGACGCTGGGATTGGAGGGCCAGTTTCCCGTGGAAGTTCGCTACGAAACCCCGATATTCGGGGCAAACGTAACTCCATTGAGGTGAAGCAAGAAAACGAACAAAACCCGATCGAGTTTGTTTTAGAAAGCTTCCTCAACTCAAACGAAACCTCCAAACCCAGGATCCATGAGAGCAGAGTTGAAGACATCCAGTTCACAGGACCCCCTATTAGCAGCATTGAGCCGACAATCACATTTGCATCAACAGAGGAGAATCACAACGACACAGTGAAAGCTATGATGCAATACTTAACCGAGACCAACCAAATGTGTCAAGAAAACAGTGTTCCCGAGCAAATAATCTTCAGCGACTTAAACAGTGTTTCCGCATCCGATTCTCTTCCAAGCGTTGAAGAATTGCTTCAAATTCCGAACGAAAAAAGCCGCAGTTTTAAACCTGACCACACTGCGAAGCACCAGCCGACGCTGAACCGCCCAAAACATTTCCACAACCAAGTAGTCAACCAGCCACATGTACCAATCCTAAACCCAGAGTTACAAAGCTTTGAAAACTTCAGCCCCGTGATAAACAACAGTCTCATCACTCCAACCAACACGTTCACATCCAACCGCATGATGGAGCTATCACCACTATCAGATCTTCATAGTTTATCGCAGGATGAAGACATGGACACAAAGATGTCGCATTATCACCATACAAGCCACCCAAACGGCCATCAATGTTTAGTATGGGCATGCAAAGCGTGTAAACGTAAAACTGGGCCACACGAtCGtCGGAGGGCGGCAACACTACGAGAGAGACGACGCCTTAAACGTGTCAACCAAGCGTACGACGCCCTTAAGCGTTGCGCATGCGCAAACCCGAACCAGAGACTTCCGAAAGTTGAGATTCTTCGCAACGCGATAACCTACATATACAATCTACAACATATGTTGTATGGCGACCAACAGTCTGATGCAAAATCACCAGAAACCAAACCAGAAACGACTTTAAGCTTGGGCGAAACTTTCGTTAGCAAAACCGAAGTTGATTCGCCGTTTTATCAGTCCGATGACGTCAGACTTACGTCATCAAGGACGTCAAGCCCGGTTGAGTCCCTTCTTGAGTCCACGTCATCGTCCTTCATAATGTCGGATTTGGGCGACGAAAATACTCAACCTCAGGTACTTTAA

>Ciona MRF (transcript variant 2, with sgRNA mismatches) START STOP

ATGACGTGTATCTCTCTAGAGGAGCTCGACCTCTCTTCAATATTCTCCAACAGCAGCAGCTATTTCACCAGTTACGCCACAAACCCCATTATGACGTCACAAAAACGGACGCCTTCACGACTGAAGCGAGCAAGCAGCGACGTTTTGTTGTCAGATACAGCGGTTTCGCCGGTAAGTCGAGAAGTTTCGGGGGTTTTATCGGAACTGGACGAACTGAAGAGATGtGTcGAGGGAAATTATATCGGAATCGATGCGGAAAAGAGCGATATCTCGATACTTGAGGAACTTAGTAATATGGCGTCCGGTTGTAGCGATTCAGACGTCGCTTACTCGTCCCCGGATTCGAGGTACGGGTCAACTGGTAACCTTACCAGCAAGACAAGTTTCGGATCGGGAATGGGTTTCCGTGACGCTGGGATTGGAGGGCCAGTTTCCCGTGGAAGTTCGCTACGAAACCCCGATATTCGGGGCAAACGTAACTCCATTGAGGTGAAGCAAGAAAACGAACAAAACCCGATCGAGTTTGTTTTAGAAAGCTTCCTCAACTCAAACGAAACCTCCAAACCCAGGATCCATGAGAGCAGAGTTGAAGACATCCAGTTCACAGGACCCCCTATTAGCAGCATTGAGCCGACAATCACATTTGCATCAACAGAGGAGAATCACAACGACACAGTGAAAGCTATGATGCAATACTTAACCGAGACCAACCAAATGTGTCAAGAAAACAGTGTTCCCGAGCAAATAATCTTCAGCGACTTAAACAGTGTTTCCGCATCCGATTCTCTTCCAAGCGTTGAAGAATTGCTTCAAATTCCGAACGAAAAAAGCCGCAGTTTTAAACCTGACCACACTGCGAAGCACCAGCCGACGCTGAACCGCCCAAAACATTTCCACAACCAAGTAGTCAACCAGCCACATGTACCAATCCTAAACCCAGAGTTACAAAGCTTTGAAAACTTCAGCCCCGTGATAAACAACAGTCTCATCACTCCAACCAACACGTTCACATCCAACCGCATGATGGAGCTATCACCACTATCAGATCTTCATAGTTTATCGCAGGATGAAGACATGGACACAAAGATGTCGCATTATCACCATACAAGCCACCCAAACGGCCATCAATGTTTAGTATGGGCATGCAAAGCGTGTAAACGTAAAACTGGGCCACACGAtCGtCGGAGGGCGGCAACACTACGAGAGAGACGACGCCTTAAACGTGTCAACCAAGCGTACGACGCCCTTAAGCGTTGCGCATGCGCAAACCCGAACCAGAGACTTCCGAAAGTTGAGATTCTTCGCAACGCGATAACCTACATATACAATCTACAACATATGTTGTATGGCGACCAACAGTCTGATGCAAAATCACCAGAAACCAAACCAGAAACGACTTTAAGCTTGGGCGAAACTTTCGTTAGCAAAACCGAAGTTGATTCGCCGTTTTATCAGTCCGATGACGTCAGACTTACGTCATCAAGGACGTCAAGCCCGGTTGAGTCCCTTCTTGAGTCCACGTCATCGTCCTTCATAATGTCGGATTTGGGCGACGAAAATACTCAACCTCAGGATACGTTCCCGGTAAACTTGCTTACTGATGACGTCACCAGACCCTCGTCAACAACCCCTGACGTCATCGCTGTCGTAAACGAACCAACAACAACAACGGAAGATCGAAACTCGAGCCCTGTGACGTCAGTGAGCAACAGCGACACCAAGGGAGCTTCTAGTTTGGTTTGTTTGACGTCGATCGTGGAAAGGATCGATTAA

>Human MYOD1 (codon optimized for C. elegans) START STOP

ATGGAACTGTTATCACCGCCTCTTCGAGATGTCGATCTTACAGCACCAGATGGTTCCCTGTGTAGTTTCGCAACTACAGATGACTTTTACGATGATCCATGCTTTGATTCTCCAGATCTTCGTTTCTTTGAGGATCTCGATCCAAGATTGATGCATGTCGGTGCTTTGTTAAAGCCTGAGGAACATTCACATTTTCCAGCAGCTGTCCATCCTGCTCCAGGAGCTCGCGAAGATGAACACGTACGGGCACCATCAGGACATCATCAAGCTGGTCGATGTCTTCTTTGGGCTTGTAAAGCTTGTAAACGAAAGACAACGAATGCAGATCGTAGAAAAGCTGCTACAATGAGAGAAAGAAGAAGACTTAGTAAGGTGAACGAAGCTTTCGAGACGCTTAAACGTTGTACTTCAAGCAACCCTAATCAAAGGCTCCCAAAAGTTGAAATTTTAAGAAATGCAATTAGATACATAGAAGGATTGCAAGCCCTACTTCGTGATCAAGATGCTGCACCGCCGGGAGCAGCTGCAGCATTTTACGCCCCAGGACCATTGCCACCAGGACGAGGTGGAGAACATTATTCAGGGGATTCAGATGCTTCATCACCCCGAAGCAATTGTTCGGATGGAATGATGGATTATTCTGGTCCGCCCTCAGGGGCTAGAAGACGAAATTGTTATGAGGGTGCTTATTATAATGAAGCCCCGTCTGAGCCTAGACCTGGCAAATCAGCAGCTGTCAGTTCATTGGATTGTTTAAGTAGTATTGTTGAACGAATTAGCACAGAATCGCCAGCCGCACCAGCTCTACTTTTAGCAGATGTCCCGAGTGAAAGTCCACCCCGGAGACAGGAAGCAGCAGCTCCAAGTGAAGGTGAATCGAGTGGTGATCCAACACAATCCCCTGATGCTGCACCACAATGTCCCGCTGGAGCTAATCCAAATCCAATTTATCAAGTTTTGTAG

>Ciona Mymk promoter (-508/-1), mEbf

tgctctggaaaatttaccaagggaaactccctcacgtggtaacatagaaatagtacaaaaaatgcttaaaattgtttttcaccaaaactaacggtacacataccgttgcctaaactaggcaaaacacatattagattgcatttaaactggtaaatgtcctaataatttaaaacttgcattaacaataaatttagttgccaataatataacttttaaattgcgtattttaagtatgtacatagttaagaatatttattcgggccataaatgtgtattttaaagttgtttatatttagaccaaatactgatattttaataaacgttttgggtcgtgagaaatatcacattattgtgtccagctgaatcagacaacagctgaagaccacatgccaattAAAaagataatggtgcagcacgcgcaaaacaaccgagagtaccgtagtacatttggtatgtcgcctttagagcattcattgcatagtgttaaaccgtagattaaatcatagtt

>Ciona Mymk promoter (-508/-1), mMRF-136

tgctctggaaaatttaccaagggaaactccctcacgtggtaacatagaaatagtacaaaaaatgcttaaaattgtttttcaccaaaactaacggtacacataccgttgcctaaactaggcaaaacacatattagattgcatttaaactggtaaatgtcctaataatttaaaacttgcattaacaataaatttagttgccaataatataacttttaaattgcgtattttaagtatgtacatagttaagaatatttattcgggccataaatgtgtattttaaagttgtttatatttagaccaaatactgatattttaataaacgttttgggtcgtgagaaatatcacattattgtgtccagctgaatcagacaaACgcGTaagaccacatgccaattcccaagataatggtgcagcacgcgcaaaacaaccgagagtaccgtagtacatttggtatgtcgcctttagagcattcattgcatagtgttaaaccgtagattaaatcatagtt

>Ciona Mymk promoter (-508/-1), mMRF-152

tgctctggaaaatttaccaagggaaactccctcacgtggtaacatagaaatagtacaaaaaatgcttaaaattgtttttcaccaaaactaacggtacacataccgttgcctaaactaggcaaaacacatattagattgcatttaaactggtaaatgtcctaataatttaaaacttgcattaacaataaatttagttgccaataatataacttttaaattgcgtattttaagtatgtacatagttaagaatatttattcgggccataaatgtgtattttaaagttgtttatatttagaccaaatactgatattttaataaacgttttgggtcgtgagaaatatcacattattgtgtcACgcGTaatcagacaacagctgaagaccacatgccaattcccaagataatggtgcagcacgcgcaaaacaaccgagagtaccgtagtacatttggtatgtcgcctttagagcattcattgcatagtgttaaaccgtagattaaatcatagtt

> Ciona Mymk promoter (-508/-1), mHES

tgctctggaaaatttaccaagggaaactccctACcgGTgtaacatagaaatagtacaaaaaatgcttaaaattgtttttcaccaaaactaacggtacacataccgttgcctaaactaggcaaaacacatattagattgcatttaaactggtaaatgtcctaataatttaaaacttgcattaacaataaatttagttgccaataatataacttttaaattgcgtattttaagtatgtacatagttaagaatatttattcgggccataaatgtgtattttaaagttgtttatatttagaccaaatactgatattttaataaacgttttgggtcgtgagaaatatcacattattgtgtccagctgaatcagacaacagctgaagaccacatgccaattcccaagataatggtgcagcacgcgcaaaacaaccgagagtaccgtagtacatttggtatgtcgcctttagagcattcattgcatagtgttaaaccgtagattaaatcatagtt

>Human MYMK -427/+3, based on Zhang et al. 2020 START

tctctactaaaaatacaaaaattagctgggtgtggtggcaggcacctgtcatcccacctattccagaggctaaggcaggagaatctcttgaacctggaaggtggaggttgcagtgagccgagatcacaccactgcactccagcctgggtgacagggcgagactctgtctcagaaaaagaaaagaaaaatcagagacaaatgctggtcacgtggcatgtcagctgttggccctctccgtgtgttactcgacacgtgcacgcacatccccgcctccgtcaagggcatttaaaccctcttgtgggtgctccccgcagctgccatcagagccctgcccaaagggagctggccttcccacttcgtgctcctgtgctggggacctgggacaccagcaccctccccaccccagccagtgctttcctcctggcccatg

>Chicken Mymk -499/+3, based on Luo et al. 2015

tgctcctattcctccctgcagagggaaagctcagctcctgaatttctatggggttgagtttcacccttcctgagcacccaggcacacgcgtgtataggtggcatttcacaccggataggatgctggctcagatttgacgctgcttccatgtctgttgctccccagcacctctcctccaggaataaaagccagattcagagctccgtgggtgtgtgcaataaacgcgtgtgcatcacagccagcatgaagcgaaccacagccgtcaaaggacacaaacagccggggcattgcagggcacggagcatttgctgccccaagaattttgctatccaagaaggaatggagcacatgggtggctcagctgtgccactgcctgcagctgttccctgcctgcagctctccgggcatttaaaatttaaatccttttctccacgtctgttttctcctccatctccagcagctcacacggacccaggagccgccggccctccaggcaccc

>Ciona Mymk -508/-1 optEbf

tgctctggaaaatttaccaagggaaactccctcacgtggtaacatagaaatagtacaaaaaatgcttaaaattgtttttcaccaaaactaacggtacacataccgttgcctaaactaggcaaaacacatattagattgcatttaaactggtaaatgtcctaataatttaaaacttgcattaacaataaatttagttgccaataatataacttttaaattgcgtattttaagtatgtacatagttaagaatatttattcgggccataaatgtgtattttaaagttgtttatatttagaccaaatactgatattttaataaacgttttgggtcgtgagaaatatcacattattgtgtccagctgaatcagacaacagctgaagaccacatgccaaCCcccaGgGGaatggtgcagcacgcgcaaaacaaccgagagtaccgtagtacatttggtatgtcgcctttagagcattcattgcatagtgttaaaccgtagattaaatcatagtt

>Ciona Mymk -508/-1 optMRF-136

tgctctggaaaatttaccaagggaaactccctcacgtggtaacatagaaatagtacaaaaaatgcttaaaattgtttttcaccaaaactaacggtacacataccgttgcctaaactaggcaaaacacatattagattgcatttaaactggtaaatgtcctaataatttaaaacttgcattaacaataaatttagttgccaataatataacttttaaattgcgtattttaagtatgtacatagttaagaatatttattcgggccataaatgtgtattttaaagttgtttatatttagaccaaatactgatattttaataaacgttttgggtcgtgagaaatatcacattattgtgtccagctgaatcagacGGcagctgCCgaccacatgccaattcccaagataatggtgcagcacgcgcaaaacaaccgagagtaccgtagtacatttggtatgtcgcctttagagcattcattgcatagtgttaaaccgtagattaaatcatagtt

>Ciona Mymk -508/-1 optMRF-152

tgctctggaaaatttaccaagggaaactccctcacgtggtaacatagaaatagtacaaaaaatgcttaaaattgtttttcaccaaaactaacggtacacataccgttgcctaaactaggcaaaacacatattagattgcatttaaactggtaaatgtcctaataatttaaaacttgcattaacaataaatttagttgccaataatataacttttaaattgcgtattttaagtatgtacatagttaagaatatttattcgggccataaatgtgtattttaaagttgtttatatttagaccaaatactgatattttaataaacgttttgggtcgtgagaaatatcacattattgtgGGcagctgCCtcagacaacagctgaagaccacatgccaattcccaagataatggtgcagcacgcgcaaaacaaccgagagtaccgtagtacatttggtatgtcgcctttagagcattcattgcatagtgttaaaccgtagattaaatcatagtt

>Ciona Mymk -508/-1 optMRF-152 + optMRF-136

tgctctggaaaatttaccaagggaaactccctcacgtggtaacatagaaatagtacaaaaaatgcttaaaattgtttttcaccaaaactaacggtacacataccgttgcctaaactaggcaaaacacatattagattgcatttaaactggtaaatgtcctaataatttaaaacttgcattaacaataaatttagttgccaataatataacttttaaattgcgtattttaagtatgtacatagttaagaatatttattcgggccataaatgtgtattttaaagttgtttatatttagaccaaatactgatattttaataaacgttttgggtcgtgagaaatatcacattattgtgGGcagctgCCtcagacGGcagctgCCgaccacatgccaattcccaagataatggtgcagcacgcgcaaaacaaccgagagtaccgtagtacatttggtatgtcgcctttagagcattcattgcatagtgttaaaccgtagattaaatcatagtt

>Ciona Mymk -508/-1 optMRF-152 + optMRF-136 + mEbf

tgctctggaaaatttaccaagggaaactccctcacgtggtaacatagaaatagtacaaaaaatgcttaaaattgtttttcaccaaaactaacggtacacataccgttgcctaaactaggcaaaacacatattagattgcatttaaactggtaaatgtcctaataatttaaaacttgcattaacaataaatttagttgccaataatataacttttaaattgcgtattttaagtatgtacatagttaagaatatttattcgggccataaatgtgtattttaaagttgtttatatttagaccaaatactgatattttaataaacgttttgggtcgtgagaaatatcacattattgtgGGcagctgCCtcagacGGcagctgCCgaccacatgccattAAAaagataatggtgcagcacgcgcaaaacaaccgagagtaccgtagtacatttggtatgtcgcctttagagcattcattgcatagtgttaaaccgtagattaaatcatagtt

**sgRNAs:**

MRF.2 sgRNA **G**+(**N19**)

**GGACGAACTGAAGAGATGCG**

MRF.3 sgRNA **G**+(**N19**)

**GTAGTGTTGCCGCCCTCCGT**

Control sgRNA **G**+(**N19**) from Stolfi et al. 2014

**GCTTTGCTACGATCTACATT**

**Electroporation mixes:**

MRF CRISPR in ASM lineage

100 ug Mymk>Unc-76::GFP

40 ug U6>MRF.2

40 ug U6>MRF.3

40 ug Mesp>Cas9

40 ug Mesp>mScarlet

Negative control CRISPR in ASM lineage

100 ug Mymk>Unc-76::GFP

80 ug U6>Control

40 ug Mesp>Cas9

40 ug Mesp>mScarlet

MRF>Ebf to see ectopic Mymk reporter in tail muscles

50 ug MRF>Ebf

70 ug Mymk>Unc-76::GFP

10 ug MRF>H2B::mCherry

MRF>EBF3(human) to see ectopic Mymk reporter in tail muscles

50 ug MRF>EBF3(human)

70 ug Mymk>Unc-76::GFP

10 ug MRF>H2B::mCherry

Negative control to compare to Ebf/EBF3 overexpression

70 ug Mymk>Unc-76::GFP

10 ug MRF>H2B::mCherry

Ebf>MRF(Tv1) to see ectopic Mymk reporter in larval CNS

50 ug Ebf>MRF(Tv1)

70 ug Mymk>Unc-76::GFP

50 ug Ebf>CD4::mCherry

Ebf>MRF(Tv2) to see ectopic Mymk reporter in larval CNS

50 ug Ebf>MRF(Tv1)

70 ug Mymk>Unc-76::GFP

50 ug Ebf>CD4::mCherry

Ebf>MYOD1(human) to see ectopic Mymk reporter in larval CNS

50 ug Ebf>MYOD1(human)

70 ug Mymk>Unc-76::GFP

50 ug Ebf>CD4::mCherry

Negative control to compare to MRF/MYOD1 overexpression

50 ug Ebf>lacZ

70 ug Mymk>Unc-76::GFP

50 ug Ebf>CD4::mCherry

MRF>MRF(Tv1) to test effect of increased MRF dose on Mymk reporter

50 ug MRF>MRF(Tv1)

70 ug Mymk>Unc-76::GFP

10 ug MRF>H2B::mCherry

MRF>Ebf to see tail muscle cell shape at 24 hpf

50 ug MRF>Ebf

30 ug MRF>CD4::GFP

10 ug MRF>H2B::mCherry

Negative control to see tail muscle cell shape at 24 hpf

30 ug MRF>CD4::GFP

10 ug MRF>H2B::mCherry

MRF>Ebf for RNAseq

50 ug MRF>Ebf

Negative control for RNAseq

50 ug MRF>CD4::GFP

Wild type Mymk reporter scoring at 44 hpf

100 ug Mymk(wt)>Unc-76::GFP

100 ug Mymk(wt)>Unc-76::mCherry

Mymk (mEbf) reporter scoring at 44 hpf

100 ug Mymk(mEbf)>Unc-76::GFP

100 ug Mymk(wt)>Unc-76::mCherry

Mymk (mMRF-136) reporter scoring at 44 hpf

100 ug Mymk(mMRF-136)>Unc-76::GFP

100 ug Mymk(wt)>Unc-76::mCherry

Mymk (mMRF-136+mEbf) reporter scoring at 44 hpf

100 ug Mymk(mMRF-136+mEbf)>Unc-76::GFP

100 ug Mymk(wt)>Unc-76::mCherry

Mymk (mMRF-152) reporter scoring at 44 hpf

100 ug Mymk(mMRF-152)>Unc-76::GFP

100 ug Mymk(wt)>Unc-76::mCherry

Mymk (mHES) reporter scoring at 24 or 44 hpf

100 ug Mymk(mHES)>Unc-76::GFP

Wild type Mymk to compare to mHES reporter at 24 or 44 hpf

100 ug Mymk(wt)>Unc-76::GFP

Human MYMK -427/+3 reporter test

100 ug Human MYMK -427/+3>Unc-76::GFP

Chicken Mymk -499/-1 reporter test

100 ug Chicken Mymk -499/-1>Unc-76::GFP

Scoring wild type Mymk reporter expression in larval CNS

50 ug Ebf>lacZ

70 ug Mymk(wt)>Unc-76::GFP

50 ug Ebf>CD4::mCherry

Scoring effect of optimized Ebf site on expression in larval CNS

70 ug Mymk(optEbf)>Unc-76::GFP

50 ug Ebf>CD4::mCherry

Scoring wild type Mymk reporter expression in tail muscles

70 ug Mymk(wt)>Unc-76::GFP

10 ug MRF>H2B::mCherry

Scoring effect of optimized MRF-152 site on expression in tail muscles

70 ug Mymk(optMRF-152)>Unc-76::GFP

10 ug MRF>H2B::mCherry

Scoring effect of optimized MRF-136 site on expression in tail muscles

70 ug Mymk(optMRF-136)>Unc-76::GFP

10 ug MRF>H2B::mCherry

Expression in tail muscles/neurons with optimized MRF-152/-136 sites

70 ug Mymk(optMRF-152 + optMRF-136)>Unc-76::GFP

10 ug MRF>H2B::mCherry

Scoring effect of optimized MRF sites + mEbf on ASM expression

100 ug Mymk(optMRF-152 + optMRF-136 + mEbf)>Unc-76::GFP

100 ug Mymk(wt)>Unc-76::mCherry

Scoring wild type Mymk reporter expression to compare to optMRFs+mEbf

100 ug Mymk(wt)>Unc-76::GFP

100 ug Mymk(wt)>Unc-76::mCherry

**Protein sequences (related to Figure 2A):**

>Ciona EBF (KH.L24.10.v1.A.SL1-1)
MATMAGPQLSGPAVRGWMQTTLVEPMPNGNVGLHRAHFEKQPPNNLRKSNFFHFVLALYDRQGQPVEIERSAFVGFVENETEIAGEKTNNGIQYRLQLLYHSGVRTEQDVFVRLIDSATKQSITYEGQDKNPEMRRVLLTHEIMCSRCCDKKSCGNRNETPSDPVVIDRYFLKFFLKCNQNCLKNAGNPRDMRRFQVVVSTTVHVDGHVLAVSDNMFVHNNSKHGRRARRVDPSEASPTIKAINPAEGWTTGGATVVIVGENFFDGLQVVFGSMVVWSELITQHAIRVQTPPRHLPGVVEVTLSYKNKQFCSGAPGRFVYTALNEPTLDYGFQRLLKTVPRHPGDPERLPKEIILKRAADVMEAVISRQYAPPSQMPPSAGITPPAPHLAAAPCAPPGSFVPQSASAAMAVAMNGYAAAAVSSQFGGTPDRFDTGSDSGYSRGNSVSPRNGYSPQTTPHSLNSGSIGSMVGLTTVGAVPAPAPYHCAPSFNSYSSASGPTFTNMNNTSPGLFSGSGIIPPSPHNGMNPLPSSGTTPGIFSFSPANMISAAKQKSAFAPVHRPHNSPSPLAPSNGNIALNGYS

>Ciona MRF KH.C14.307.v1.A.SL1-1 (Transcript Variant 2)

MTCISLEELDLSSIFSNSSSYFTSYATNPIMTSQKRTPSRLKRASSDVLLSDTAVSPVSREVSGVLSELDELKRCVEGNYIGIDAEKSDISILEELSNMASGCSDSDVAYSSPDSRYGSTGNLTSKTSFGSGMGFRDAGIGGPVSRGSSLRNPDIRGKRNSIEVKQENEQNPIEFVLESFLNSNETSKPRIHESRVEDIQFTGPPISSIEPTITFASTEENHNDTVKAMMQYLTETNQMCQENSVPEQIIFSDLNSVSASDSLPSVEELLQIPNEKSRSFKPDHTAKHQPTLNRPKHFHNQVVNQPHVPILNPELQSFENFSPVINNSLITPTNTFTSNRMMELSPLSDLHSLSQDEDMDTKMSHYHHTSHPNGHQCLVWACKACKRKTGPHDRRRAATLRERRRLKRVNQAYDALKRCACANPNQRLPKVEILRNAITYIYNLQHMLYGDQQSDAKSPETKPETTLSLGETFVSKTEVDSPFYQSDDVRLTSSRTSSPVESLLESTSSSFIMSDLGDENTQPQDTFPVNLLTDDVTRPSSTTPDVIAVVNEPTTTTEDRNSSPVTSVSNSDTKGASSLVCLTSIVERID

>Human MYOD1 (CAA40000.1)

MELLSPPLRDVDLTAPDGSLCSFATTDDFYDDPCFDSPDLRFFEDLDPRLMHVGALLKPEEHSHFPAAVH

PAPGAREDEHVRAPSGHHQAGRCLLWACKACKRKTTNADRRKAATMRERRRLSKVNEAFETLKRCTSSNP

NQRLPKVEILRNAIRYIEGLQALLRDQDAAPPGAAAFYAPGPLPPGRGGEHYSGDSDASSPRSNCSDGMM

DYSGPPSGARRRNCYEGAYYNEAPSEPRPGKSAAVSSLDYLSSIVERISTESPAAPALLLADVPSESPPR

RQEAAAPSEGESSGDPTQSPDAAPQCPAGANPNPIYQVL

**qPCR primers (relates to Figure 2B):**

| **Gene name** | **Strand** | **Sequence (5’ to 3’)** | **Amplicon size** |
| --- | --- | --- | --- |
| Human MYMK | Forward | CTTCATGCGTCACGACATCCT | 103bp |
|  | Reverse | CCTCTTGGGTTCGTCGAAGT |  |
| Human 18S | Forward | GTAACCCGTTGAACCCCATT | 151bp |
|  | Reverse | CCATCCAATCGGTAGTAGCG |  |
